# Functional modules predict genetic interactions, revealing evolutionary conservation and rewiring

**DOI:** 10.64898/2025.12.18.695201

**Authors:** Chenchu Lin, Veronica Gheorghe, Juihsuan Chou, Sabriyeh Alibai, Subin Kim, Nazanin Esmaeili Anvar, Yixin Xu, Xingdi Ma, Lori L. Wilson, Russell D. Moser, Christopher J. Kemp, Junjie Chen, Scott Kopetz, Traver Hart

**Affiliations:** Department of Systems Biology, The University of Texas MD Anderson Cancer Center, Houston, TX, USA; MD Anderson/UT Health Graduate School of Biomedical Sciences, Houston, TX, USA; Department of Gastrointestinal Medical Oncology, The University of Texas MD Anderson Cancer Center, Houston, TX USA; Division of Human Biology, Fred Hutchinson Cancer Research Center, Seattle, WA, USA; Department of Experimental Radiation Oncology, The University of Texas MD Anderson Cancer Center, Houston, TX, USA

## Abstract

Genetic interactions can reveal gene function and identify disease-relevant synthetic lethals, but resource-intensive screening in human cells constrains experimental scope and limits our understanding of how generalizable sparse hits are. Here, we leverage principles from yeast genetic networks to identify human gene modules involved in receptor tyrosine kinase signaling and DNA damage response that are predicted to be enriched for genetic interactions, then use our enCas12a-based In4mer combinatorial knockout platform to efficiently screen these modules across diverse cancer cell lines. We identify hundreds of unreported synthetic lethals, including a dense network within the protein N-linked glycosylation machinery that shows both evolutionary conservation and network rewiring, and we confirm that interactions discovered in 2D cell culture are maintained in more physiologically relevant models. Critically, a meta-analysis within and between screens highlights sources of variation in published results, reveals the potential for false positives in screens with many essential genes, and demonstrates the need to move beyond DDR for actionable synthetic lethalities.

## Main

Genetic interactions provide a useful framework for deciphering how genes work together to control the emergence of phenotype at the cellular and organismal level. Genetic interaction occurs when the combined effect of perturbing two genes deviates from the expected effect from single perturbation^1,2^. Synthetic lethality, the limiting case of genetic interaction where double knockout causes cell death, has long offered a compelling paradigm for cancer^3^, through an array of related but distinct exploits. For example, identification of vulnerabilities arising from tumor-specific genetic lesions grew from the discovery of PARP inhibitor sensitivity in BRCA-mutated cancers^4,5^, later expanding to include homologous recombination deficiency more broadly. More recently, tumors with defective mismatch repair machinery were discovered to be specifically sensitive to loss of Werner helicase (WRN)^6,7^.

Extending this approach beyond tumor-specific lesions to identify, e.g., independently druggable SL gene pairs with synergistic activity (Fig. 1A) requires combinatorial perturbation approaches. In the budding yeast *Saccharomyces cerevisiae,* knockouts of over 23 million double mutants identified hundreds of thousands of quantitative genetic interactions yielding a global network that covers more than 90% of yeast genes^8,9^. This comprehensive network established several foundational principles: genes encoding proteins that function together in the same pathway or complex tend to share similar patterns of genetic interactions; these correlated genetic interaction profiles define functional modules corresponding to protein complexes, biological processes, and biochemical pathways; and genetic interactions are enriched within and between functionally related modules^10–12^.

**Figure 1.**
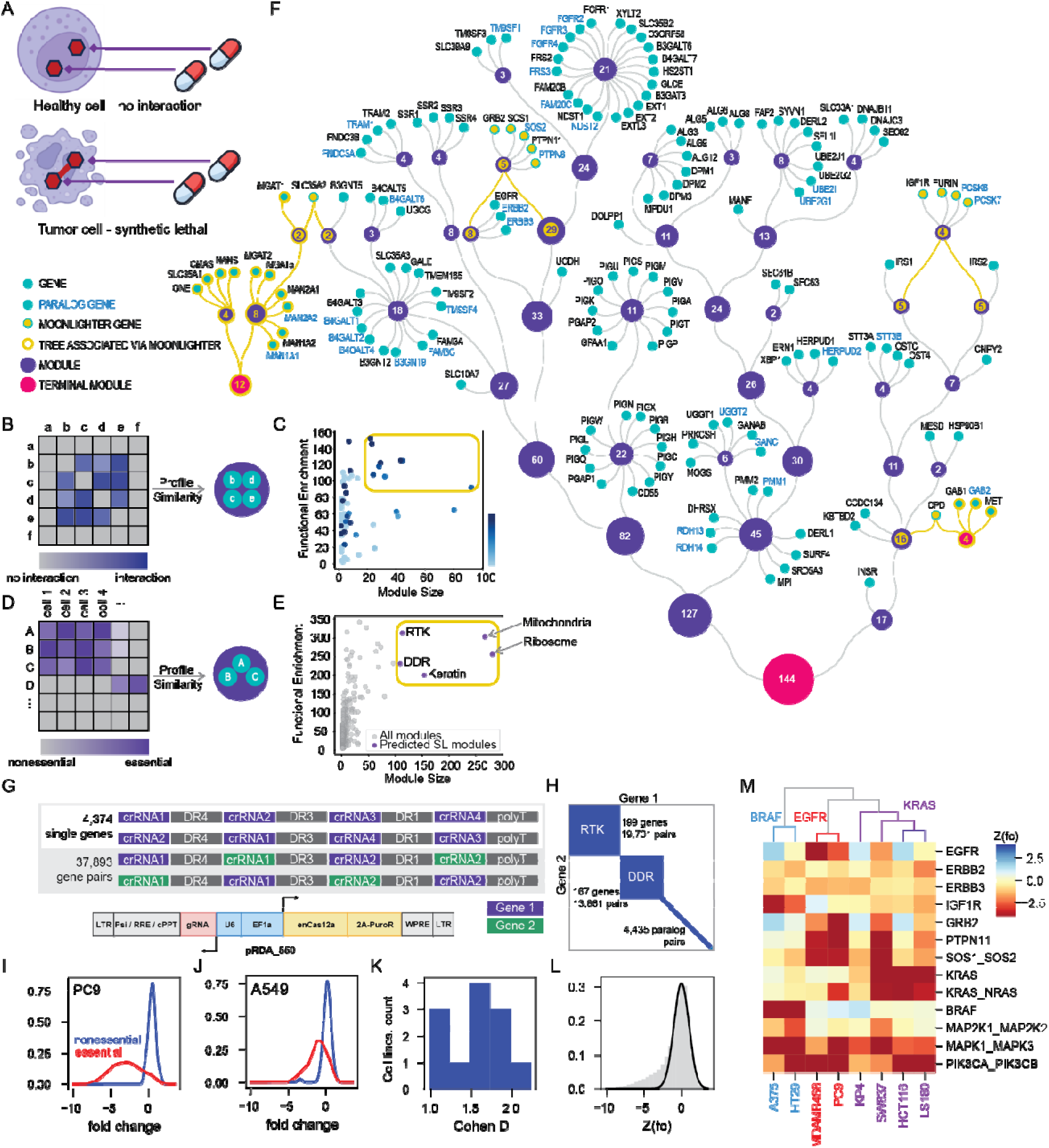
Network-guided design of genetic interaction screens. (A) Goals of this study. Identifying and targeting of tumor-specific synthetic lethals is a goal for precision medicine. (B) Yeast genetic interaction networks of double knockouts of gene 1 (x-axis) and gene 2 (y-axis), where profile similarity of genetic interaction profiles indicates similar biological function, creating a functional interaction network. (C) Dissecting the yeast functional network suggests that large modules (x-axis) with high functional coherence (y-axis) are enriched for genetic interactions (colorbar), providing a rubric for prioritizing genetic interaction screens. (D) CRISPR screens in cancer cell lines provide a similar insight into functional relationships between genes: correlated fitness profiles indicate functional similarity. (E) Decomposition of the human functional interaction network identifies large modules (x-axis) with high functional coherence (y-axis), that we hypothesize are enriched for genetic interactions. (F) The RTK module as constructed using our network decomposition approach whereby genes (black label) are progressively collapsed into modules. After the structure is learned, paralogs (blue label) are added to each module. Numbers on each module (purple) indicate number of genes. Moonlighters (yellow) are associated with two different modules; see Methods for details. (G) Design of Cas12a guide arrays targeting single and double gene knockouts, using In4mer constructs. (H) A single library targets all pairs of 199 RTK genes, all pairs of 167 DDR genes, and all individual 4,435 paralog pairs. 104 paralogs are duplicated in the RTK and DDR modules. (I-J) Kernel density estimates of guide arrays targeting control essential (red) and nonessential (blue) genes in a high-quality screen, PC9, and a lower quality screen, A549. (K) Separation of essentials and nonessentials was used to measure Cohen’s D for each cell line. Eight screens with Cohen’s D > 1.5 were deemed high quality. (L) For each screen, fold changes across all targets were normalized to a Z-score by fitting the main peak of the fold change distribution to a Gaussian. (M) Clustering of Zfc across 13 targets in the RTK/MAP kinase cascade in eight high-quality cell lines clearly distinguishes *BRAF*^mut^, *KRAS*^mut^, and EGFR-dependent cells.

Replicating yeast genetic interaction networks in human cells is considerably more challenging due to the larger genome, greater biological complexity, and cell-type heterogeneity. Genome-wide CRISPR/Cas9 knockout screening across hundreds of cell lines paved the way for discovery of context-specific genetic vulnerabilities as well as the construction of co-essentiality networks^6,13–16^ that indicate functional relationships. Subsequently, the development of combinatorial and multiplex perturbation technologies enabled direct measurement of genetic interaction in human cells^17–29^, but at high experimental cost. Large-scale screen efforts by Horlbeck *et al.*^21^ and recently by Fielden *et al.*^20^ used over 1 million and nearly 680,000 dual-guide Cas9 constructs, respectively, with each replicate requiring about half a billion cells. Though highly informative, these studies each assayed only ∼0.1% of the possible search space in one or two cell lines, highlighting the resource-intensiveness and technical challenge of combinatorial screening in mammalian cells. Considering the cost, scale and limited reproducibility of large-scale screens, some studies have focused on exploration of paralogs^30–32^, which often exhibit functional redundancy and are more enriched for synthetic lethality. Together, these limitations highlight the need for improved multiplex technologies and predictive models that prioritize genes with high interaction potential.

Here, we leverage principles from the architecture of the yeast genetic interaction network to predict which regions of the human genome are likely to be enriched for genetic interactions, then test the top predictions with our CRISPR/Cas12a In4mer platform. In4mer uses the RNA-processing capability of the Cas12a CRISPR endonuclease to process multiple guide RNAs expressed from a single transcript, enabling simultaneous targeting of multiple genes, each with multiple guides^33,34^. We and others have shown that enAsCas12a shows comparable or better efficiency than Cas9 platforms for surveying genetic interactions^33,35–37^, while reducing the number of constructs required to assay GI by as much as five-fold^36^. Our screens identify hundreds of synthetic lethal and suppressor interactions, including extensive interactions within the ER-localized protein glycosylation machinery that show strong evidence of both evolutionary conservation and rewiring. We show that interactions discovered in permissive 2D culture are also observed in more physiologically relevant environments including 3D organoid models and mouse xenografts. We confirm that our network-guided predictions yield substantial enrichment of genetic interactions, establishing a principled framework for prioritizing combinatorial perturbation experiments in human cells.

### Network-guided prediction of genetic interactions

In yeast screens, correlated genetic interaction profiles imply co-functionality (Fig. 1B), giving rise to functional interaction networks that both predict gene function and identify modules of coherent biological activity. It has previously been observed that these functional modules tend to be enriched for genetic interactions between their constituent genes, both in yeast^10–12^ and recently in human cells^26,38^, and, conversely, that interactions between genes in unrelated modules are extremely rare^8,9^. We built on this knowledge to develop a method for decomposition of functional interaction networks that would facilitate discovery of modules with high proportion of genetic interactions (Supplementary Fig. 1A-E). We discovered that, for the yeast network, the combination of relatively large module size and high functional coherence within the module combine to nominate specific gene sets that were enriched for within-module genetic interactions (Fig. 1C). We then took human gene essentiality data from CRISPR screens in cancer cell lines^39,40^, constructed an optimized functional interaction network using methods we have described previously^41^ (Fig. 1D), and decomposed the human network using the same principles. We identified five large modules with very high functional enrichment (Fig. 1E), three of which (mitochondria, ribosome, and keratin) are relatively homogeneous clusters with low biological variation. We focused on the remaining two modules for further study: one containing receptor tyrosine kinase (RTK) genes and their biogenesis factors, and another with DNA damage response (DDR) genes.

The RTK module (Fig. 1F) is characterized by oncogenic receptors that show high variation in gene essentiality across DepMap cell lines, including *EGFR, FGFR1, IGF1R, MET*, and *INSR*. Our functional network decomposition approach includes “moonlighter” genes and modules that are connected to disparate other modules in a context-dependent manner; e.g. the *GRB2/SOS1/PTPN11* signal transduction module is connected with both the *EGFR* and *FGFR1* modules. Close paralogs of each gene are added to each module; e.g. *SOS1* paralog *SOS2* is added to the signal transduction module. Notably, this global RTK network includes functional submodules encoding ER-based protein glycosylation and ER-associated degradation, and *FGFR1*-associated heparin sulfate biosynthesis, all elements required for the proper posttranslational processing and trafficking of cell surface receptors^42,43^. However, this module did not contain the classical downstream signaling components such as the MAP kinase and PI3 kinase pathways, where mutations disconnect them from upstream signaling nodes^14^; we manually added these clusters where appropriate (Supplementary Fig. 1F). The total RTK module expanded from 144 genes in the main cluster to 199 genes across all clusters that we predicted would be enriched for genetic interactions.

To test these predictions, we developed a CRISPR genetic interaction library based on our In4mer platform^36^. In4mer is a Cas12a-based system that expresses arrays of four individually addressable guide RNA. Our library design includes two arrays targeting each single gene with four guides each, and two arrays targeting each gene pair with two guides targeting each gene (Fig. 1G). The single library targets all combinations of 199 genes in the RTK module, all combinations of 167 genes in the DDR module, as well as all 4,435 paralog pairs from our prior catalog, along with controls, in a single pool of 85k sequences (Fig. 1H; Supplementary table S1-S4).

We screened 12 adherent carcinoma cell lines with the library and evaluated each for quality control (Supplementary Figs. 2, 3; Supplementary table S5-S7). Separation of essential and nonessential genes ranged from excellent (PC9 cells, Fig. 1I) to moderate (A549 cells, Fig. 1J). Based on Cohen’s D statistics across the 12 cell lines, which showed a bimodal distribution of effect sizes (Fig. 1K), we selected the top eight (Cohen’s D >= 1.5) for detailed analysis. To facilitate comparison across cell lines, we normalized each screen’s fold change to a Z score (Zfc) by fitting the main peak of the overall fold change distribution to a normal distribution (Fig. 1L, Supplementary table S8). Clustering of selected marker genes or gene pairs using Zfc clearly differentiated the eight high-quality screens by their oncogenic driver genes (Fig. 1M). Notably, *BRAF^mut^*cells (A375, HT29) also show strong dependence on *IGF1R* while *EGFR* cells (MDA-MB-468 and PC9) show strong dependence on the *GRB2/SOS1/PTPN11* module. *KRAS*^mut^ cells are generally independent of upstream signaling elements, though SW837 cells are dependent on both *KRAS* and *EGFR* signaling. Overall, these preliminary results indicate the CRISPR screens are working as designed and provide a clean view of the genetic dependencies of each cell line.

### Strong, context-specific genetic interactions in the RTK module

To decipher genetic interactions from combinatorial knockouts, we developed the Genetic interaction Regression Analysis of Pairwise Effects (GRAPE) algorithm^44^ (Fig. 2A). GRAPE builds a regression model with a binary predictor matrix and learns the single gene knockout phenotypes (beta coefficients) that best fit the observed double gene knockout data. Raw genetic interactions are then measured as the difference between observed and expected fold change and are converted to Z-scores using a sliding window to estimate local variance, which increases with the severity of single knockout phenotype. As an example, the observed fold changes for all pairwise knockouts for the PC9 cell line are compared with the fold changes expected from the sum of the beta coefficients for each gene pair (Fig. 2B; Supplementary Figs. 4, 5). Raw genetic interaction scores as well as a local standard deviation are calculated (Fig. 2C), and Z scores of genetic interaction (Zgi, Fig. 2D; Supplementary Fig. 6) show synthetic lethal interactions among paralog pairs in the RTK and DDR libraries. We also estimated a normalized effect size for each interaction, where an effect size of 1 is achieved when all guide arrays targeting the gene pair have zero reads across all replicates of a screen (see Methods; Supplementary Fig. 7), and effect size zero is where observed and expected fold change are equal (Fig. 2E; Supplementary Fig. 8; Supplementary table S9). We defined a high-effect-size (HE) hit as FDR < 0.25 from Zgi and Effect size > 0.15, with an additional requirement for strong double knockout phenotype (Zfc < –2). We previously showed that using both genetic interaction and fold change thresholds is important for identifying high-quality paralog synthetic lethals^45^.

**Figure 2.**
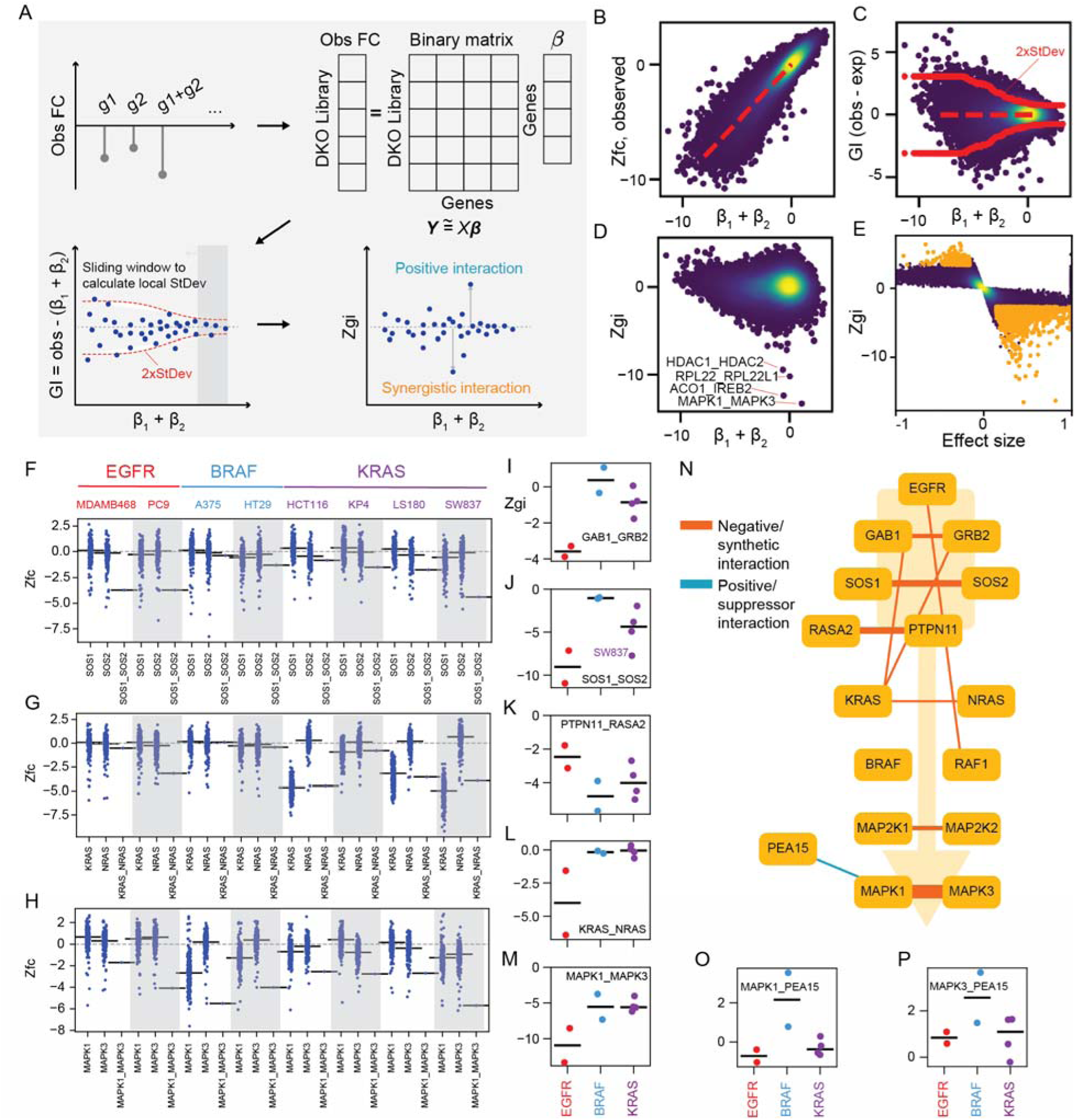
Global and background-specific genetic interactions. (A) The GRAPE algorithm learns regression coefficients that estimate single gene knockout phenotype (fold change), calculates an expected double knockout phenotype, measures the deviation from this phenotype, and then calculates a Z-score of the deviation. (B) Observed Zfc (y-axis) vs. Zfc predicted from single gene regression coefficients (x-axis). (C) Raw GI score is observed – expected, as a function of expected Zfc. A sliding window estimates local standard deviation (red) for calculating Z-score of GI. (D) Z score of GI (y-axis) vs. predicted Zfc (x-axis). Negative GI are synthetic lethal; labeled pairs are synthetic lethal paralog pairs. (E) Zgi (y-axis) vs. normalized effect size (x-axis) for each gene pair. Orange indicates pairs with ES > 0.15, Zgi FDR < 0.25. (F) Raw Zfc for the SOS1_SOS2 paralog pair across all 8 high quality screens. SOS1: Zfc of all genes paired with SOS1. SOS2: Zfc of all genes paired with SOS2. Black segment indicates beta score (expected single knockout Zfc) from GRAPE. (G-H) Zfc plots for background-specific KRAS_NRAS and global MAPK1_MAPK3 synthetic lethality. (I-M) Negative/synthetic Zgi scores for labeled gene pairs in each cell line, separated by oncogenic driver (red, EGFR; blue, BRAF; purple, KRAS). (N) RTK and MAP kinase signaling network, showing strong synthetic and suppressor interactions. (O-P) Positive/suppressor Zgi scores for MAPK1/MAPK3_PEA15 in each cell line, separated by oncogenic driver.

Focusing first on the RTK module, we observed strong, background-specific genetic interactions between paralogs in the RTK/MAPK signal transduction pathway. Fold change of all ∼200 gene pairs of either *SOS1* or *SOS2*, paralogous guanine nucleotide exchange factors for RAS proteins, are distributed around the beta scores for single knockouts. Double knockout shows severe fitness phenotype (Zfc ∼ –4), specifically in EGFR-dependent cells, but not in KRAS– or BRAF-mutant backgrounds (Fig. 2F), demonstrating a clear context-specific synthetic lethality. This is consistent with the established role of *SOS1* and *SOS2* as the primary RAS GEFs required for efficient GTP loading for wildtype signaling downstream of EGFR^46^. In contrast, oncogenic *KRAS* and *BRAF* are not dependent on SOS function, explaining the absence of a synthetic interaction in those genetic contexts. Prior studies have shown that loss of *SOS1* or *SOS2* impairs signaling in *EGFR*-mutant cells^47–50^; our data extends these findings by demonstrating functional redundancy between *SOS1* and *SOS2* in *EGFR*-driven models.

Similarly, RAS paralogs *KRAS* and *NRAS* show no single knockout fitness defect in *EGFR*-dependent cells, but double knockouts indicate clear synthetic lethality in *EGFR*-dependent PC9 cells (Fig. 2G). This is similar to the *KRAS-NRAS* synthetic lethality we previously observed in *BCR-ABL* fusion K562 leukemia cells^36^, but neither single nor double knockouts were lethal in MDA-MB-468 cells. This differential sensitivity may be explained by compensation from *HRAS* in MDA-MB-468 cells or may simply be a false negative from low knockout efficiency. Neither single nor double knockout of *KRAS* and *NRAS* shows phenotype in *BRAF^mut^*cells, consistent with *BRAF* functioning downstream of RAS, while the severe double knockout phenotype in *KRAS^mut^* cells is entirely explained by the *KRAS* single knockout (Fig. 2G), an example of mutation-driven epistasis. ERK genes *MAPK1* and *MAPK3*, paralogs at the end of the MAP kinase cascade, show strong synthetic lethality across all eight cells (Fig. 2H).

Moving beyond paralogs, a summary view illustrates which genetic interactions are background specific. Taking the Zgi score for each gene pair and performing a linear regression across the three genotypes identifies *GAB1/GRB2* and *SOS1/SOS2* as synthetic lethal in the EGFR background, consistent with these genes’ roles in the RTK signal transduction complex (Figs. 2I, J). In contrast, *PTPN11/RASA2* and *MAPK1/3*, while strong across all three backgrounds, show relative weakness (*PTPN11/RASA2*) and strength (*KRAS/NRAS, MAPK1/3*) in EGFR relative to the RAS/RAF mutants (Figs. 2K-N).

Suppressor interactions are also present. Loss of *PEA15* reduces the single-gene dependency on either *MAPK1* or *MAPK3* in *BRAF*-mutant cells (Figs. 2N-P). *PEA15* encodes a protein that sequesters activated MAP kinase protein in the cytoplasm, and *BRAF^V600E^*melanoma cells are generally associated with a strong preference for *MAPK1* over *MAPK3*^13,41,51^. Loss of *PEA15* leads to increased nuclear translocation of MAP kinase protein^52–54^, likely offsetting any loss of proliferation signaling by ERK in its transcriptional activation role when one copy is lost to genetic perturbation. In the A375 melanoma cells in our panel, which are highly dependent on *MAPK1* (GRAPE beta score –5.3), the *MAPK1/PEA15* suppressor interaction is a stringent hit (FDR 0.21, Effect size –0.427) while the *MAPK3/PEA15* interaction is just below threshold (FDR 0.17, Effect size –0.148). Overall, this subnetwork shows both known and novel genetic interactions, and highlights the dominance of paralog synthetic lethality in genetic interaction networks.

### Integration across screens to improve sensitivity of detection

Across the eight high-quality cell lines, we find a median of 77 HE interactions per screen, with a strong relationship between screen quality and number of SL hits discovered (Spearman ρ = – 0.86, P=0.007; Fig. 3A). However, only 8 synthetic lethal gene pairs in the RTK library are observed in at least half of the screens (Fig. 3B). The total number of unique HE hits across all 8 screens (n=331) represents a small fraction of the total number of gene pairs assayed in the RTK library (1.7%), with each screen recovering a median of 52 hits (0.3% of tested pairs). Given the low hit rate in individual cell lines, we sought to integrate data across the eight screens to attempt to separate consistent but weaker interactions from background noise. Importantly, the per-cell-line Zgi score is normally distributed for each independent cell line (Fig. 3C), providing a robust statistical foundation for our integrative approach. We calculated an integrated Z-score for each gene pair across all eight cell lines (hereafter Zint), using Strube’s method to correct for P-value inflation from correlated samples (see Supplementary Methods; Supplementary table S10). The resulting Zint shows a similar dynamic range, retains its normal distribution (Fig. 3D), and yields 300 SL at FDR < 0.25 (0.9% of tested pairs).

**Figure 3.**
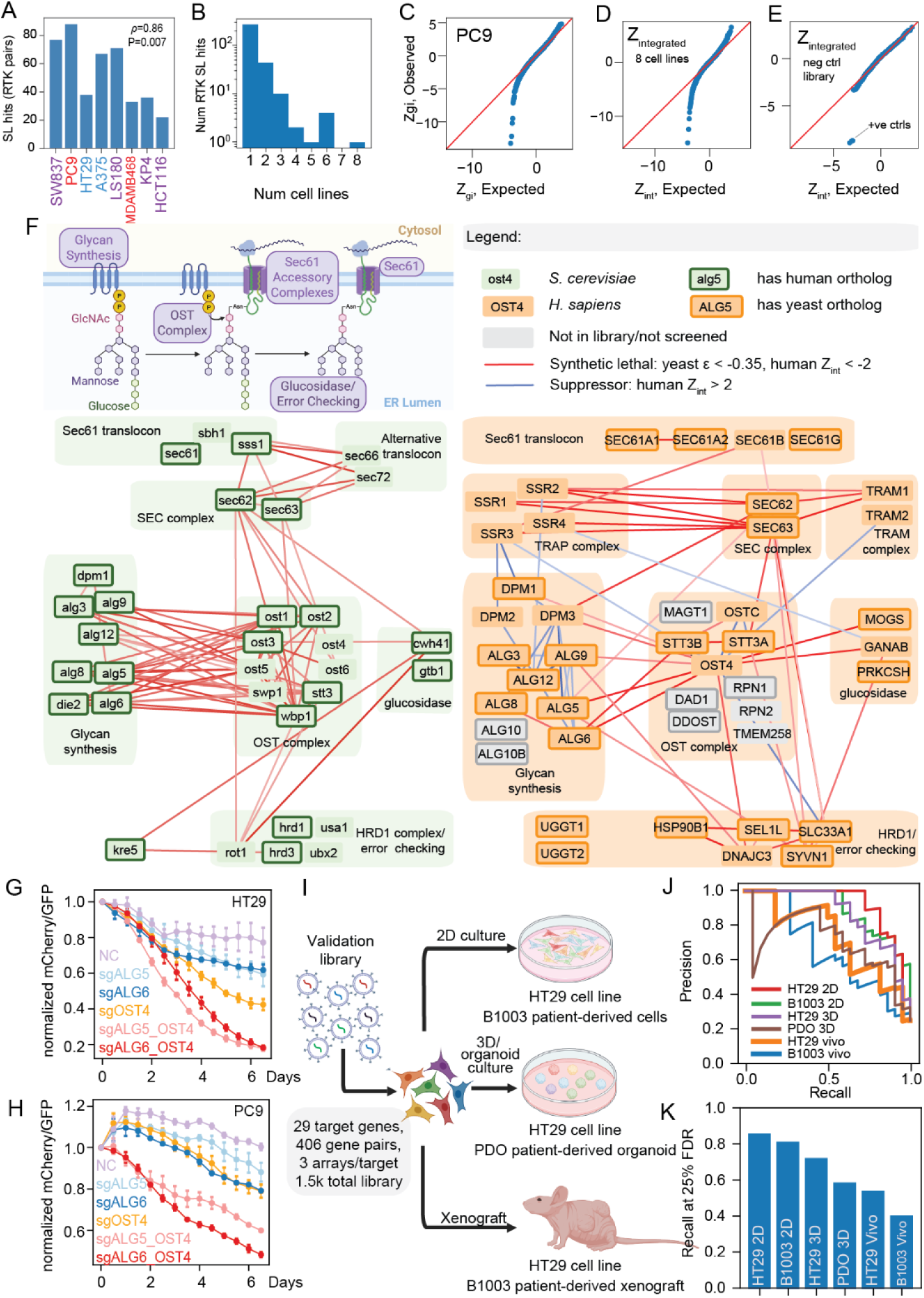
Integrative analysis of interactions in the RTK network. (A) Number of high effect size (HE) negative interactions across eight high-quality screens in RTK module. Inset: Spearman correlation between screen quality and number of HE hits. (B) Distribution of HE negative interaction hits across eight high-quality screens in RTK module. (C) Q-Q plot of Zgi for a single screen. Diagonal indicates normal distribution. (D) Q-Q plot of integrated Z-scores across all eight high-quality screens, with similar dynamic range with single-screen distribution. (E) Q-Q plot of integrated Z-scores in negative control library screens across 3 cell lines. Only positive controls pairs are significant. (F) Protein glycosylation pathway schematic (top left) and corresponding genetic interaction subnetworks in yeast (bottom left) and human cells (bottom right). (G-H) Color competition assays for two novel interactions, ALG5_OST4 and ALG6_OST4, in HT29 (G) and PC9 (H) cells. (I) A 1.5k validation library for testing whether interactions in 2D are also present in more complex models. (J) Precision-recall plots for six validation library screens. (K) Recall at 25% FDR (75% precision) for each of the six validation screens confirms interactions with high effect size are present in 3D and *in vivo* models.

To estimate the background density of genetic interactions, we randomly selected 64 genes with a broad range of single knockout phenotypes but no meaningful coessentiality in DepMap. We added three positive control paralog pairs (six genes), for a total of 70 genes. We constructed a genetic interaction library targeting all pairs of these genes with three arrays targeting each gene pair and screened the resulting 6k library in three cell lines (Supplementary Fig. 9), filtering targets with low T0 read counts. We applied the GRAPE analytical pipeline and found that only the positive control paralogs met the threshold for high effect size hits. We integrated Zgi scores for 1,915 gene pairs across the three screens and observed that, beyond the three positive controls, there are zero hits at FDR < 0.25 (the top hit has Benjamini-adjusted P-value 0.265), and all remaining Zint scores are normally distributed (Fig. 3E; Supplementary table S11). We conclude that the background frequency of genetic interactions observed using our perturbation technology and informatics pipeline is well under 1 in 1,000, that our pipeline effectively samples the null distribution of observed genetic interaction scores, and that our FDR thresholds are likely conservative.

Mapping integrated interactions from the eight high-quality RTK screens onto the hierarchical network from Figure 1 identifies a dense network in the ER-localized N-linked protein glycosylation pathway (Fig. 3F) that shows similar network architecture from yeast to human. The highly conserved N-linked glycosylation process involves a glycan synthesis pathway, encoded by the ALG (“asparagine-linked glycosylation”) genes, that constructs the glycan tree. The OST complex, which occurs in mammals as two isoforms with a shared *OST4* subunit, is co-located with the Sec61 complex on the ER membrane and transfers the completed glycan onto the translocating peptide chain at specific asparagine residues^55,56^. The Sec61 translocon channel is itself composed of a core complex and a series of accessory complexes, including the SEC (*SEC62/SEC63*) complex conserved throughout eukaryotic evolution, and the TRAP and TRAM complexes that are also widely conserved but have been lost in laboratory Saccharomyces strains. The three glucose residues that terminate the glycan tree serve as an error-checking mechanism that is “read” by the *HRD1* complex, with nascent polypeptides failing this check being shunted to the ER-associated degradation (ERAD) pathway^57^.

Examining yeast genetic interactions^8,9^ within this highly conserved molecular machinery reveals a dense network of synthetic lethals between components of this module. In particular, most members of the glycan synthesis pathway are synthetic lethal with most members of the OST complex (Fig. 3F). This is notable because we display only the strongest synthetic lethals from the yeast data (yeast genetic interaction score, epsilon, < –0.35; background frequency ∼ 0.1% of all gene pairs across the dataset), indicating that this is one of the densest concentrations of synthetic lethals across the entire yeast network.

Mapping our interactions among these same protein complexes and pathways (Fig. 3F) shows strong indication of both conservation and rewiring of the genetic interaction network. The mammalian Sec61 channel is encoded by two paralogs, *SEC61A1* and *SEC61A2*, which are not in our coessentiality-derived module but are strongly synthetic lethal in our paralog data (see below). The TRAP, SEC, and TRAM accessory complexes are all strongly synthetic lethal with each other. While none of these genes are individually essential (Supplementary Fig. 10), this synthetic lethality network suggests that any one of these complexes is dispensable but the remaining two are required. Indeed it was previously shown that when either *SEC62* or *SEC63* is lost, cells become more dependent on TRAP to stabilize Sec61 and maintain minimal translocation activity^58^. This functional redundancy, distributed across three protein subcomplexes, may explain the historical difficulty in determining the substrate specificity of each complex^59–62^.

As in yeast, we observe strong synthetic lethality within the OST complex across our screens. *OST4* is required for efficient N-glycosylation by stabilizing the two native mammalian OST complexes^63^. These complex isoforms are distinguished by paralogous catalytic subunits *STT3A/B*, among the strongest synthetic lethals across all our screens, with the *STT3A* complex isoform showing positive interactions between its subunits *OSTC-STT3A* and *OSTC-OST4*. Also conserved are the interactions between OST and the glycan synthesis genes, especially *OST4-ALG5,* which catalyzes the formation of the donor-substrate dolichol phosphate-glucose, and *OST4-ALG6,* which transfers the first of three terminal glucose residues onto the oligosaccharide. Absence of the terminal glucose impairs glycan recognition and leads to ERAD. We also observe suppressor interactions between members of the glycan synthase pathway, consistent with epistatic buffering of a linear pathway. Collectively, the discovery of an evolutionarily conserved genetic interaction cluster, connected with another that has been rewired by the disappearance of protein complexes, strongly supports the rigor of our network-guided predictive model coupled with In4mer-based genetic interaction screening.

### Interactions maintained in physiologically relevant models

A fundamental question about genetic screens in cell lines is whether the genetic dependencies and interactions discovered in 2D culture hold any relevance in more physiologically relevant conditions. We validated several strong novel interactions *in vitro*, with color competition assays with *OST4* knockout paired with either *ALG5* or *ALG6* showing rapid depletion in both HT29 and PC9 cells (Figs. 3G, H). These interactions are evident even after short incubations, consistent with high effect size and high penetrance. To test whether these are maintained in other growth conditions, we constructed a small validation library for testing high-effect hits in different models (Fig. 3I). The library probes all pairs of 29 target genes, primarily drawn from the genes shown in the subnetwork in Figure 3F (Supplementary table S12). We screened the library in the B1003 patient-derived colorectal tumor cells in 2D culture and in mouse xenografts, in a patient-derived organoid model PDO065 grown in 3D culture, and in HT29 cells grown in 2D and 3D cultures as well as in mouse xenografts (Supplementary table S13, S14). We found that the 2D cultures outperformed 3D and *in vivo* (Figs. 3J, K; Supplementary Fig. 11), but that the highest effect size hits were still detectable even using mouse xenograft models, with both HT29 and B1003 xenograft screens achieving ∼50% recall of top hits at 25% FDR. Thus, robust hits in 2D models appear to be good predictors of genetic interactions in more physiologically relevant contexts, while the lower sensitivity of *in vivo* models clearly argues for initial screening *in vitro*.

### Paralog synthetic lethals across eight cancer cell lines

Paralogs are a rich source of synthetic lethals compared to unrelated gene pairs, which not only boosts their scientific and potentially clinical relevance but also provides a useful sandbox for technology development. Several labs, including ours, have performed paralog synthetic lethality screens^30–32,35,37^ using digenic perturbation modalities with an array of Cas9 and Cas12a endonucleases and guide expression constructs. A common informatics approach in these projects is to measure paralog synthetic lethality by delta log fold change (dLFC), directly comparing double knockout phenotype with single knockout phenotype (Fig. 4A). We have previously shown that Z-normalizing across all observed dLFC values in a screen (Fig. 4B) leads to greater consistency of hit calls across different screens^45^, a finding that supports the normalizing approaches in some paralog screens^30,31^. This methodology highlights a fundamental difference between paralog screens and broader genetic interaction screens, in that paralog screens rely on a much smaller measurement pool for calculating single gene knockout phenotype and, by extension, expected double knockout phenotype. In our RTK genetic interaction pools, we intentionally expanded the genelist to include close paralogs of nodes in the network module, resulting in 89 paralog pairs across eight cell lines (n=712 total observations) being tested using both the GRAPE and ZdLFC pipelines. Overall, there is very strong concordance between the two approaches (Fig. 4C), unless one gene shows a strong single knockout phenotype. In these cases, GRAPE corrects for the variance-phenotype relationship (Fig. 2A, C), but ZdLFC does not carry this information. We judge GRAPE results to be more accurate, since their single gene knockout phenotypes are estimated from substantially more data. We therefore define stringent synthetic lethals as previously described (ZdLFC < –3, double knockout Zfc < –2) with an additional robustness filter removing pairs where one gene shows a strong single knockout (SKO) phenotype (Zfc < –3.5).

**Figure 4.**
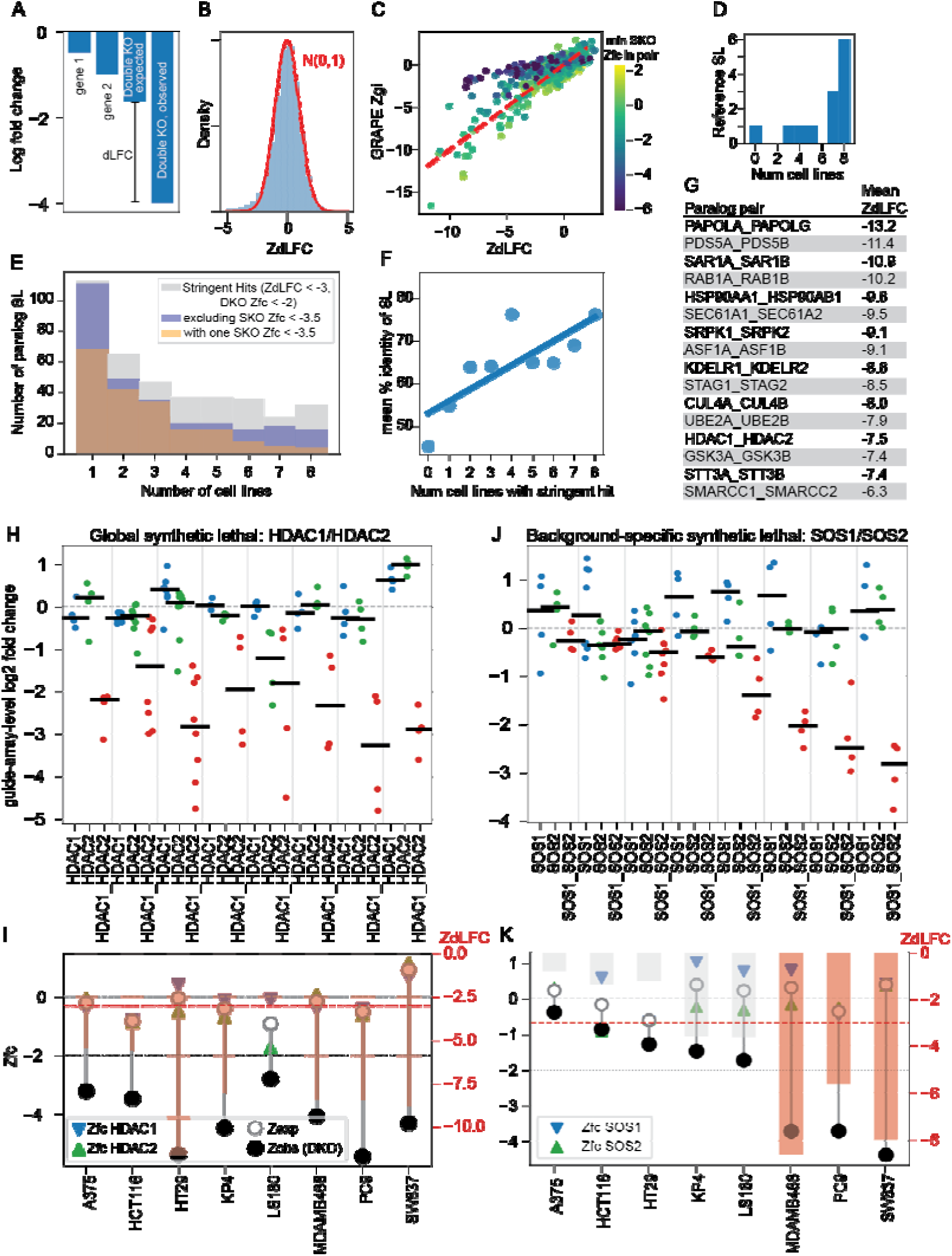
Integrative analysis of paralog genetic interactions. (A) Delta log fold change (dLFC) is a commonly used approach for quantifying paralog synthetic lethality. (B) Z-transformed dLFC (ZdLFC) is a robust scoring for identifying synthetic lethal interactions in paralog screens. (C) Comparison of GRAPE Zgi and ZdLFC across paralogs in genetic interaction modules, colored by most severe single knockout phenotype in pair. (D) Distribution of reference SL pairs according to the number of eight high-quality screens in which they were identified as hits. (E) Number of paralog SL hits across eight high-quality screens by different hit-calling threshold. (F) Correlation between paralog SL hits and protein sequence identity. (G) Top SL paralog pairs ranked by mean ZdLFC across eight cell lines. (H-I) HDAC1-HDAC2 as an example of global synthetic lethal pairs. (H) Guide-array-level LFC for HDAC1 and HDAC2 single and pairwise perturbations in each cell line. (I) Zfc (left y-axis) and the difference between expected and observed pairwise effects across cell lines. Green and blue indicate HDAC1 and HDAC2 single knockouts, respectively, and open and closed circles represent expected and observed Zfc. Overlaid with bar chart of ZdLFC (right y-axis): red, ZdLFC < –3 threshold; gray, does not exceed threshold. (J–K) SOS1–SOS2 as an example of context-specific synthetic-lethal paralogs. (J) Guide-array-level LFC for SOS1 and SOS2 single and pairs. (K) Zfc and the difference between expected and observed pairwise effects across cell lines.

We used this approach to score the 4,435 paralog pairs in our combined library (Supplementary table S15). Nine of the 13 reference synthetic lethals from Esmaeili Anvar & Lin, *et al*.^36^ are stringent hits in at least seven of the eight screens, while one (*CSNK1D/E*) did not meet the threshold in any screen (Fig. 4D). A total of 391 unique synthetic lethal pairs are observed across the eight screens, with 106 removed by the SKO phenotype filter, resulting in 285 stringent hits. Of those, 90 (31%) are observed in at least half of the screens, while 111 (39%) are scored in a single screen (Fig. 4E). The number of cell lines in which a hit is observed is strongly correlated with the similarity of the two proteins (Fig. 4F). The top synthetic lethals, as measured by mean ZdLFC score across 8 screens (Fig. 4G), contain many of the Esmaeili Anvar & Lin, *et al.*, reference paralogs as well as the key paralogs from the RTK glycosylation network (*SEC61A1/2*, *STT3A/B*) and *HDAC1/2*, which we use as positive controls.

The *HDAC1/2* pair is a good example of a pan-synthetic lethal paralog. Single gene knockouts consistently show near zero-fold change, while double knockout shows severe phenotype across most reagents (Fig. 4H). This translates into consistent differences between expected and observed double knockout fold change (Fig. 4I), resulting in strong synthetic lethality across all eight backgrounds. In contrast, background-specific synthetic lethal *SOS1/2* shows single and double knockout phenotype in the range of (−1,1) across most cell lines (Fig. 4J), but strong double knockout phenotype in *EGFR*-dependent cell lines MDAMB468, PC9, and SW837. Weaker signal in LS180 and KP4 cells meets the ZdLFC threshold (bar chart, Fig. 4K, right y-axis), but not the double knockout Zfc threshold (scatter plot, Fig. 4K, left y-axis), and is not classified as a stringent hit.

### Genetic interactions in the DDR module

We constructed the DDR sublibrary using the same approach as the RTK library. The DDR module in our coessentiality network contains a hierarchical network with functionally coherent end nodes (Fig. 5A, highlights). We manually added other correlated (*MDM2* cluster) and anticorrelated (*TP53* cluster) smaller modules known to be involved in DDR, for a total of 167 genes (Supplementary table S4). These were included in the GRAPE analysis described for the RTK sublibrary. The DDR library yielded many more SL hits per cell line (median 190; range 152-286) than RTK, with no relationship between screen quality and number of hits (Fig. 5B). These encompass a total of 1,083 unique gene pairs, or 8% of the DDR gene pairs tested. Of those, only 62 (0.5% of tested pairs) are hits in at least half of the screens (Fig. 5C). SL hits greatly outnumber suppressor hits in the DDR screens (Figs. 5D, E). Paralogs are again among the top common HE synthetic lethals, including iron sensors *ACO1/IREB2* (n=7 screens) and ribosomal proteins (and possibly cryptic tumor suppressors) *RPL22/RPL22L1*^64^ (n=4 screens).

**Figure 5.**
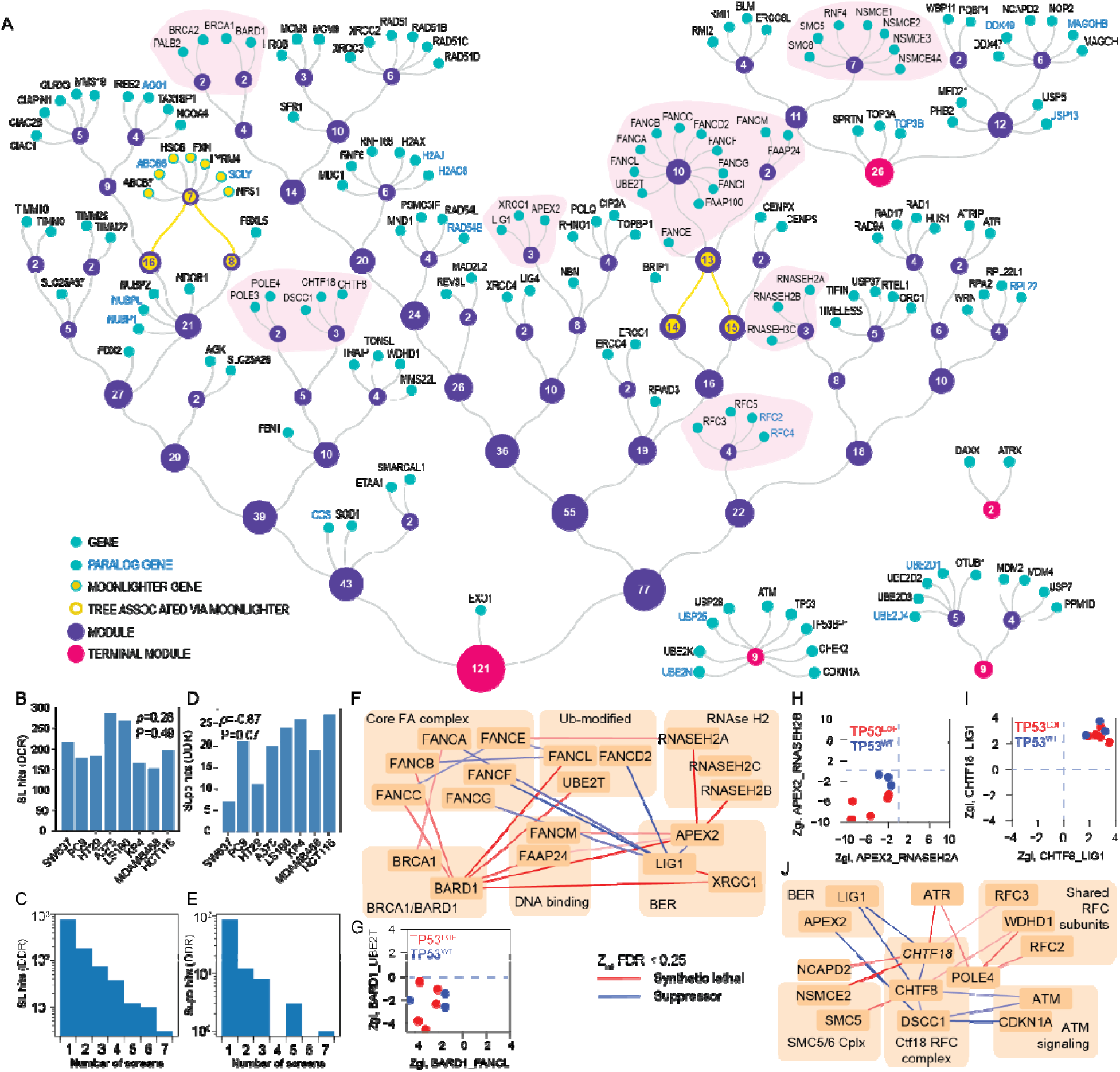
Integrative analysis of interactions in the DDR network. (A) The primary DDR module learned from decomposition of the coessentiality network. Node labels indicate module size (number of genes). Inset: manually added modules. (B) Number of HE negative interactions across eight high-quality screens in DDR module. Inset: Spearman correlation between screen quality and number of HE hits. (C) Distribution of HE negative interaction hits across eight high-quality screens in DDR module. (D) Number of HE positive interactions across eight high-quality screens in DDR module. (E) Distribution of HE positive interaction hits across eight high-quality screens in DDR module. (F) Genetic interactions (Zint FDR < 0.25) centered on the FA complex. (G) Similarity of genetic interactions between co-complex genes (FANCL, UBE2T). Zgi scores for interactions in each cell line. Color: TP53 status of cells. (H) Zgi scores for APEX2_RNASEH2A and APEX2_RNASEH2B, highlighting TP53 status-dependent variation. (I) Zgi scores for CHTF18-LIG1 and CHTF8-LIG1 pairs, labeled by TP53 status. (J) Genetic interactions (Zint FDR < 0.25) centered on the Ctf18 complex.

Using the previously calculated Zint scores to aggregate signal across screens, we mapped strong interactions onto selected functional modules (Fig. 5F). The *BRCA1/BARD1* module recapitulates its known synthetic lethal relationship with the Fanconi anemia cluster, and the integrated scores in Figure 5F for the *BARD1/FANCL* and *BARD1/UBE2T* interactions are consistent with the per-cell-line Zgi scores (Fig. 5G). Similarly, the broadly reported synthetic lethality between *APEX2* and the RNASEH2 complex^65–67^ is among the strongest hits in our data but shows a clear bias toward *TP53*^LOF^ cell lines (Fig. 5H). A similar subnetwork centered on the Ctf18-containing alternate RFC complex, which facilitates *PCNA* loading for leading strand synthesis^68–70^ (Fig. 5I), shows the classic within-complex suppressor interactions (*CHTF8, CHTF18, DSCC1*) and synthetic lethal interactions between complex isoforms (*RFC2, RFC3, WDHD1*). The strongest suppressor interactions, between the Ctf18 complex members and the *LIG1/APEX2* base excision repair machinery, are again highly consistent across individual screens (Fig. 5J).

### Global and comparative analysis indicates systematic false positives

Our DDR sublibrary is considerably smaller than recently published all-by-all genetic interaction surveys by Fielden *et al.* (548 genes)^20^ and Hayward *et al.* (233 genes)^71^ – the latter of whom used the In4mer platform and GRAPE pipeline for screening and first-order hit calling – while Herken *et al.* conducted large screens (304 genes)^28^ in which a subset were DDR genes. Despite the high-level DDR description, the four groups assayed largely disjoint set of genes (Fig. 6A) and gene pairs (Fig. 6B). Among the total of 898 genes assayed across the four groups, only 21 genes (210 gene pairs) were tested by all four. Of the 210 tested pairs, only 8 hits were reported in more than one screen, and of those, half are in the well-described BRCA1/Fanconi synthetic lethality cluster (Fig. 6C).

**Figure 6.**
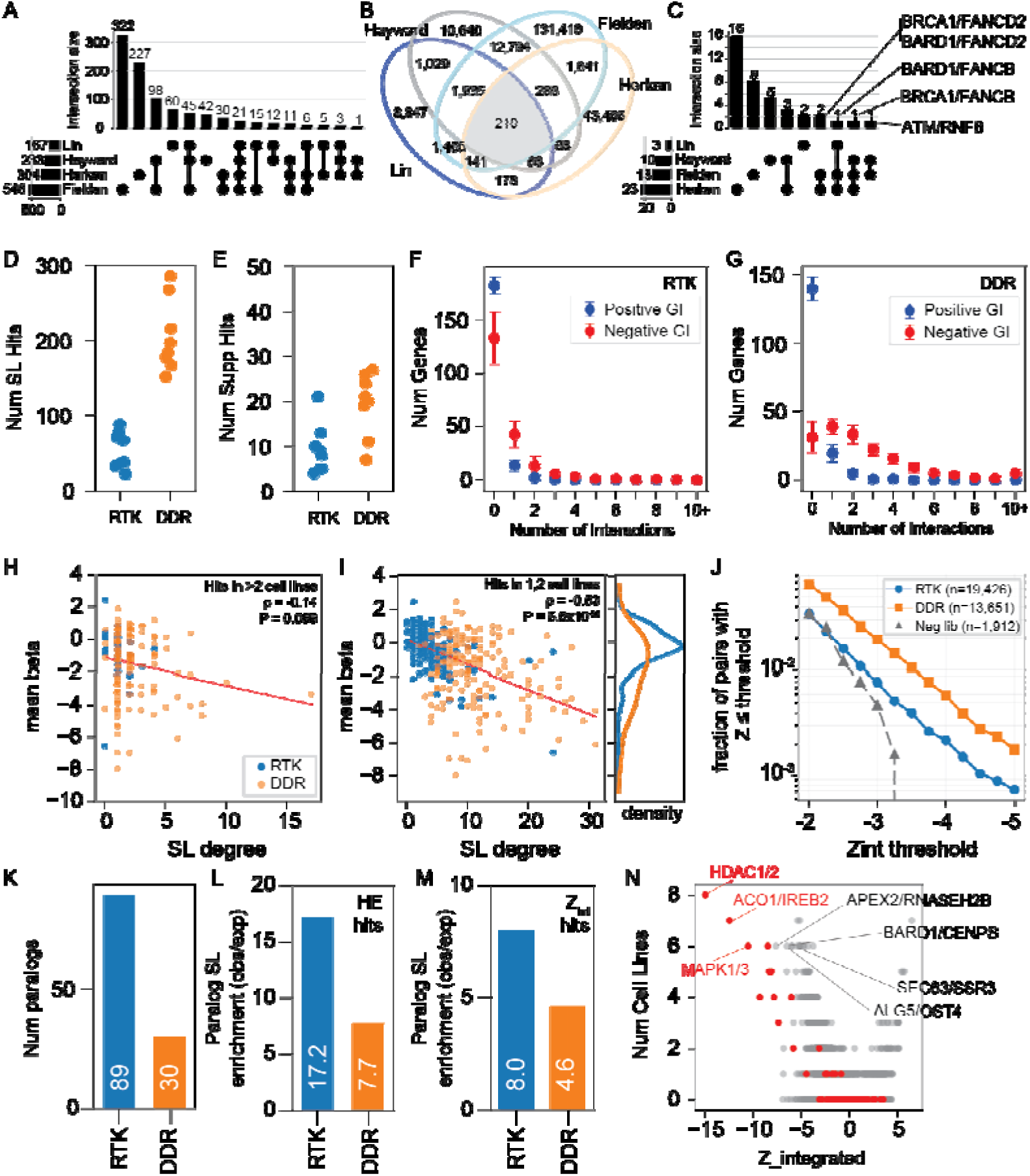
Characteristics of genetic interactions. (A) Comparison of screened gene targets among this study, Hayward *et al.*, Herken *et al.* and Fielden *et al.* (B) Overlap of gene pairs across four DDR screens. (C) Common hits identified among four DDR screens. Hits identified in more than one screen are dominated by the BRCA/Fanconi SL cluster. (D) Number of high-effect SL hits in RTK and DDR modules. Each point is one cell line. (E) Number of high-effect suppressor hits. (F) Histogram of degree distribution of negative and positive GI in the RTK module. Most genes show no interactions. Error bars are stdev across eight high-quality screens. (G) Histogram of degree distribution of negative and positive GI in the DDR module. In contrast with RTK, nearly every DDR gene has at least one high-effect SL interaction. (H-I) Correlation between single-gene knockout phenotype (mean beta) and SL degree for hits in more than two cell lines (H) or two or fewer cell lines (I), suggesting that strong single-knockout phenotypes may contribute to false-positive SL calls. (J) Using integrated (Zint) scores, fraction of total pairs (y-axis) that are called hits by threshold (x-axis). (K) Number of paralog pairs in RTK and DDR sublibraries. (L) Enrichment of paralog SL hits relative to non-paralog hits in the RTK and DDR screens. (M) Enrichment calculated using the Zint. (N) Gene pairs with Integrated Z-score (x-axis) vs. number of cell lines with a high effect size (HE) hit. Highlighted interactions are strong on both metrics, but paralogs (red) dominate the genetic interaction landscape.

This lack of overlap could be explained by (a) a large set of true interactions against which our perturbation tools have low sensitivity, resulting in false negatives; (b) a small set of true interactions against which our tools show low specificity, resulting in high false positives; or, more likely, some combination thereof. The absence of any substantial resource of gold-standard genetic interactions against which to reasonably estimate empirical false positive or false negative rates requires that we apply alternative approaches. We have elsewhere examined how different analytical pipelines can yield wildly different results when applied to the same genetic interaction screen data^44^; here we leverage our observations of two distinct screens in the same experiment across several cell lines to explore sources of bias.

First, we reiterate that the GRAPE analysis of the DDR sublibrary yields far more high-effect-size synthetic lethal hits per cell line (median 190; 1.4% of 14k pairs tested) than the RTK library (median 52 hits; 0.3% of 19k pairs tested) (Fig. 6D), while both returned a much smaller number of suppressor interactions within roughly the same range (Fig. 6E). Relatedly, most genes in the RTK network had no HE interactions in any screen (Fig. 6F), while more than half of DDR genes show two or more interactions per cell line. In both cases, positive/suppressor interactions are infrequent, and rarely does any gene have more than a single positive interaction in any cell.

Next, we consider the relationship between hit frequency (number of cell lines), SL degree (number of synthetic lethal interactions for a given gene), and the single knockout phenotype of a given gene. For gene pairs that are high-effect synthetic lethal in more than two cell lines – those in which we have the most confidence – there is no relationship between SL degree and the single knockout phenotype estimated by the GRAPE beta coefficient (Spearman ρ = –0.14, P=0.089) (Fig. 6H). In contrast, for genes with high-effect hits in only one or two cell lines, there is a strong relationship between severity of phenotype and SL degree, even beyond the phenotype-variance filter imposed by GRAPE (Spearman ρ = –0.53; P=5.6×10^−26^) (Fig. 6I). This relationship is present in the RTK genes but is dominated by the DDR genes, whose distribution of beta coefficients skews markedly more negative than the RTK genes (Fig. 6I).

From these observations, we conclude that there is a strong relationship between single knockout phenotype and false positive synthetic lethals, and that the population of DDR genes having stronger single gene knockout phenotype than the RTK genes contributes to the systematic overcalling of hits in this subnetwork. This is corroborated when considering the fraction of the library that exceeds a given Z-score threshold (Fig. 6J): RTK library hits occur at a similar rate as the negative control library in the non-significant Z-score region well-sampled by both (|Z| < 3), diverging as the Z threshold gets more stringent and true hits emerge in RTK. In contrast, the DDR library has a consistent ∼2.6-fold (range 1.9– to 3.0-fold) increase in the frequency of hits called at each threshold relative to both RTK and the random library.

While higher false discovery rates are problematic when determining the broader implications of individual interactions, including the opportunity for therapeutic exploitation, they have less impact on the functional genomics applications of genetic interactions. Billmann *et al.*^38^ use correlated genetic interaction profiles to construct a functional interaction network, and Herken^28^, Fielden^20^, and Hayward all use correlation maps to elucidate functional modules and assign gene function. This approach would be somewhat circular in our work, given that the modules themselves were defined through functional interaction networks, but rather than learn modules through clustering, we can test for enrichment of interactions between predefined clusters. Indeed, we find that synthetic lethal interactions are enriched between specific modules, including the RNase H2 complex and *BRCA1/BARD1* (Supplementary Fig. 12). Conversely, within-complex suppressor interactions are enriched in the RNase H2 complex, the 911 complex and the Ctf18 alternate RFC complex (Supplementary Fig. 13).

Finally, paralogs clearly dominate the genetic interaction landscape. Within both the DDR and RTK sublibraries, paralogs remain the primary source of high-penetrance synthetic lethal interactions with large effect sizes. There are 119 paralog pairs encoded in the two sublibraries (Fig. 6K) and, despite representing only a small fraction of gene pairs in the two modules, high effect size paralog synthetic lethals are more than 17-fold enriched compared to non-paralog SL in the RTK library, and more than 7-fold enriched among DDR gene pairs (Fig. 6L). Using the integrated Z score, which recovers more interactions across the cell lines, this enrichment is diluted (RTK, 8-fold enrichment; DDR, 4.6-fold; Fig. 6M), but the trend is maintained. Paralogs remain the strongest hits using both high-effect and Zint metrics (Fig. 6N), with the only non-paralogous genes approaching the interaction strength of paralogs being the well-described *APEX2/RNASEH2B* and *BARD1/CENPS* interactions in the DDR module and the newly discovered core interactions of the N-linked glycosylation module (Fig. 6N).

### Conclusions

Systematic mapping of genetic interactions in human cells has been limited by the immense combinatorial search space and the resource-intensive nature of multiplex perturbation screens. Here, we demonstrate that principles governing genetic interaction architecture in yeast, specifically, that functionally coherent modules are enriched for within-module interactions, can be used to prioritize regions of the human genome for combinatorial screening. By decomposing an optimized co-essentiality network, we identified a 199-gene RTK module and a 167-gene DDR module that we predicted would be enriched for within-module genetic interactions. We screened the two modules, plus more than four thousand paralog pairs, using a single In4mer-based combinatorial knockout library. With a total size of ∼85k clones, the library is comparable in size to first-generation genome-scale CRISPR/Cas9 single gene knockout screens, allowing us to screen 12 cell lines, eight of which passed our stringent quality control filters.

Our screens uncovered hundreds of genetic interactions across the two functionally distinct modules. Within the RTK signaling network, we identified both universal interactions, such as the broadly penetrant *MAPK1/MAPK3* and *STT3A/STT3B* synthetic lethalities, and context-specific vulnerabilities that depend on oncogenic background. The dense network of interactions within the ER-localized protein glycosylation machinery, connecting the glycan synthesis pathway, the oligosaccharyltransferase complex, and error-checking glucosyltransferases presents a previously unappreciated hub of synthetic lethality that is highly conserved from yeast to human. In contrast, the strong synthetic lethality between the translocon accessory complexes in human cells marks an evolutionary rewiring in response to the presence or absence of these complexes. Validation experiments demonstrated that high-effect interactions identified in 2D screens are detectable in 3D organoid cultures and mouse xenografts, with recall of top hits even in a patient-derived tumor model screened in a challenging *in vivo* setting. This suggests that robust hits from 2D screening provide a reasonable starting point for cancer target discovery, though the reduced signal-to-noise in 3D and *in vivo* models underscores the importance of prioritizing interactions with large effect sizes for translational studies.

The compact In4mer library facilitates screening in multiple cell lines, enabling evaluation of the generalizability of findings in a way that larger screens in one or two cell lines do not. One key finding is that integrating weak signal across multiple screens is a robust and sensitive method of extracting more signal while controlling error rates, which is corroborated by our discovery of an evolutionarily conserved glycosylation network. The chief drawback of this approach is that it limits the ability to identify background-specific genetic interactions.

Our second major conclusion is that genes with a significant single knockout phenotype are systematically enriched for false positive interactions. This trend is also observed in the Billman *et al*. genetic interaction study^38^, for which data was collected by conducting genome-scale Cas9 knockout screens in isogenic knockout strains of the HAP1 cell line, an entirely orthogonal approach to genetic interaction mapping. The SKO/noise relationship is largely absent in our RTK subnetwork but evident in the DDR genes. This added noise level, coupled with the absence of positive controls for genetic interactions beyond paralog synthetic lethals and a few well-described interactions (e.g. BRCA/FA), renders comparison of published DDR genetic interaction screens problematic. The top hits are almost certainly true; for example, the *FANCM/SMARCAL1* interaction characterized in the Fielden *et al.* study is among the strongest hits in our data. Beyond the top hits, however, robustness and generalizability remain open questions.

This work represents a step toward addressing these questions. We demonstrate that a functional interaction network can guide the design of a genetic interaction screen toward a target-rich geneset, within the constraints of the SKO/noise correlation. The In4mer platform enables genetic interaction screens at markedly reduced cost and effort, allowing direct measurement of generalizability and robustness by screening across multiple backgrounds. On the other hand, overall platform sensitivity has room for improvement, and the network-guided predictive model is untested for regions with high functional coherence in human cells but no corresponding density of synthetic lethals in yeast. Crucially, in the absence of gold standards against which to measure the quality of these large-scale screens, the modest overlap within our screens and between published works highlights the challenge of reproducibility in this field and suggests that both false positive and false negative rates remain significant. Within these limitations, however, the framework we establish here provides a scalable strategy for systematic genetic interaction mapping in human cells.

## Acknowledgements

We thank Drs. Tim Heffernan and Ronald DePinho (MDACC) for cell line resources. We thank Dr. Yuan-Hung Lo (MDACC) for sharing organoid screen protocols. We thank Dr. John Doench (Broad Institute) for the generous gift of the plasmid. We thank Daria Nikolaeva for the graphic design and visual presentation of the networks.

## Funding

National Institutes of Health/NCI grant U01CA275886 (TH)

National Institutes of Health/NIGMS grant R35GM130119 (TH)

National Institutes of Health/NCI grant R35CA274234 (J Chen)

NCI Cancer Center Support Grant P30CA16672

MDACC Odyssey Fellowship Program (CL)

## Author contributions

Conceptualization: CL, TH, SKo, JChe, CJK, RDM

Methodology: CL, VG, JCho, SKi, NAE

Investigation: CL, SA, SKi, YX, XM, LLW, RDM

Formal analysis: CL, VG, JCho, SKi, NAE, TH

Funding acquisition: TH, SKo, JChe, CL

Writing – original draft: CL, VG, JCho, SA, SKi, XM, LLW, TH

Writing – review and editing: all authors

## Competing interests

The authors declare no competing interests.

## Data and materials availability

All experimental data are available at the Hart lab github site: https://github.com/hart-lab/Functional_modules_predict_genetic_interactions.

## Supplementary Materials

Materials and methods, Figs S1-S13 available in Supplementary Materials file Supplementary tables S1-S16

## Notes

### Competing Interest Statement

The authors have declared no competing interest.

### Summary of Updates

This version significantly revises the analysis pipeline, adds a comparison to the yeast genetic interaction network, includes extensive new data on paralogs, and compares published genetic interaction screens in the DDR space. Figure 1 is unchanged but all subsequent figures have been revised.

