## Supplementary Methods and Figures for "Functional modules predict genetic interactions, revealing evolutionary conservation and rewiring"

### Materials and Methods

#### Network prediction of genetic interactions

Yeast functional network

Quantitative genetic interaction (GI) scores (ε) for *Saccharomyces cerevisiae* were obtained from Costanzo et al.[1], comprising ~23 million gene pairs among 5,416 genes. The ε scores quantify deviations in double-mutant fitness from multiplicative expectation, capturing both negative (synthetic-sick/lethal) and positive (suppressive) interactions. The raw matrix was cleaned and deduplicated so that each unordered gene pair was represented once. Allelic suffixes were stripped from gene identifiers, reciprocal entries were collapsed by alphabetical ordering, and the entry with the larger |ε| value was retained. Self-interactions and allele-allele comparisons were removed, yielding a non-redundant dataset of ~19 million gene pairs. Each gene's GI profile was defined as its vector of ε scores across all partners. Pairwise similarity between profiles was computed using the Pearson correlation coefficient, generating a symmetric correlation matrix used for downstream clustering and module detection.

To identify functionally coherent gene modules, we applied a hierarchical clustering algorithm based on partial correlation (see Partial correlation-based clustering below). The method iteratively merges genes or modules according to their correlation and partial correlation relationships, with a correlation cutoff ($r$) defining when merging stops. Varying $r$ controls network granularity: higher thresholds retain only very strong associations (producing small, dense clusters), whereas lower thresholds yield broader, coarser modules. Correlation thresholds were systematically scanned from $r = 0.05$ to 0.6 (step = 0.05) to evaluate the relationship between network resolution and enrichment for synthetic lethal (SL) interactions. Most merging events occurred at $r = 0.2$–0.35 (Supplementary Fig. 1A). At each threshold, the final set of non-overlapping modules was extracted for downstream enrichment analysis. Network tractability was also assessed by examining module number and size. Very loose thresholds (≤ 0.15) collapsed most genes into few large modules, whereas strict cutoffs (≥ 0.4) produced numerous tiny clusters. The threshold $r =$

0.25 yielded > 200 modules of manageable size (largest ≈ 470 genes), providing a practical balance between resolution and interpretability.

Synthetic lethality enrichment analysis

High-confidence SL interactions were defined as gene pairs with $\varepsilon < -0.35$ and $p < 0.05$, following Costanzo et al.[1], yielding 19,893 SL pairs. For each correlation threshold, we computed the following metrics using module-restricted gene pairs:

1. SL recall – fraction of SL pairs recovered within modules (both genes in the same module).
2. Coverage – fraction of experimentally tested gene pairs represented within modules, relative to the total screened pairs.
3. SL enrichment – proportion of SL pairs among tested pairs within modules, and fold-enrichment relative to the global background SL rate.

Modules with < 80% of within-pair ε scores available were excluded to prevent undersampling bias. Functional enrichment was assessed for each module using Gene Ontology (GO) annotations[2] through hypergeometric tests for over-representation, as described in the Functional enrichment analysis section below. Enrichment strength was summarized as a combined functional enrichment score of $\log_2$(fold-enrichment) × ($-\log_{10}$ adjusted *p*-value). Across correlation thresholds, SL recall, coverage, and $\log_2$ fold-enrichment were evaluated to determine the optimal network resolution (Supplementary Fig. 1B). Lower thresholds (≤ 0.2) yielded broad coverage but low specificity, while higher thresholds (≥ 0.3) increased enrichment at the cost of recall. A cutoff of $r = 0.25$ provided the best balance, achieving ~3-fold SL enrichment over background with ~10% recall and ~1% coverage of screened pairs, representing an efficient level for experimental exploration. This threshold was therefore used as the optimal resolution for defining yeast functional modules and benchmarking subsequent human network analysis.

Partial correlation-based clustering of functional networks

We constructed hierarchical functional interaction networks using an iterative partial correlation–based clustering algorithm designed to identify co-functional modules while accounting for genes with context-dependent or dual roles ("moonlighters") (Supplementary Fig. 1E). At each iteration, pairwise Pearson correlations were calculated among all current features in the matrix (genes or cluster centroids). Diagonal values were set to zero to exclude self-correlations. The algorithm proceeded through the following steps:

1. Identify the strongest pair: The feature pair with the highest Pearson correlation ($\rho_{ij}$) was selected as a candidate for merging. Iteration continued until this maximum correlation fell below a predefined threshold (defined separately for each dataset).
2. Compute first-order partial correlations: For every third feature $k$ (where $k \neq i,j$), the first-order partial correlation $\rho_{ij|k}$ was calculated using the formula:

$$\rho_{ij|k} = \frac{\rho_{ij} - \rho_{ik}\rho_{jk}}{\sqrt{(1-\rho_{ik}^2)(1-\rho_{jk}^2)}}$$

A ratio was defined as:

$$ratio_{ij|k} = \left(\frac{\rho_{ij|k}}{\rho_{ij}}\right)^2$$

This ratio quantifies how conditioning on $k$ affects the association between $i$ and $j$:

- ratio ≪ 1 → $k$ mediates the association, likely belonging to the same pathway or complex.
- ratio ≈ 1 → $k$ is independent of the pair.
- ratio ≫ 1 → conditioning increases $\rho_{ij}$, suggesting a potential moonlighting relationship.

3. Moonlighter detection: When the highest-ranked third feature exhibited $\rho_{ij|k} > \rho_{ij}$ and ratio > 1.2, we assessed the correlation signs to detect conditional connectivity:
   - If $\rho_{ik} > 0$ and $\rho_{jk} < 0$ (with both ratios > 1.2), $i$ was classified as a moonlighter.
   - If $\rho_{ik} < 0$ and $\rho_{jk} > 0$, $j$ was classified as a moonlighter.

   The identified moonlighter was duplicated into two downstream clusters, each capturing one of its positive associations. To avoid overrepresentation, its contribution to each cluster was weighted equally (half of its signal).
4. Cluster formation: For non-moonlighting cases, the selected pair ($i$,$j$) was merged with any additional features $k$ satisfying ratio < 0.6, forming a cluster. The cluster centroid was defined as the mean vector of all member profiles.
5. Matrix update and iteration: After each merge or moonlighter duplication, the corresponding columns were replaced by the new cluster centroid, and the correlation matrix was recalculated. The process was repeated until the strongest remaining correlation dropped below the predefined cutoff.

Upon termination, the algorithm produced a hierarchical set of modules and identified moonlighter genes linking multiple clusters. The final output comprised the set of clusters, their gene memberships, and the updated correlation matrix reflecting inter-cluster relationships. This procedure captures both tightly connected co-functional modules and overlapping context-dependent associations within the functional network.

Human functional network

Gene essentiality profiles were obtained from the Broad DepMap (CERES-corrected CRISPR KO data across cancer cell lines)[3,4]. The data was organized as a feature matrix of gene dependency scores. Prior to network inference, the matrix was PCA-whitened to normalize variance and remove linear dependencies, resulting in a matrix where each feature has unit variance[5]. The partial correlation–based clustering procedure was applied to the whitened matrix to construct a hierarchical functional interaction network. To extend the yeast insights to human cells, we used the yeast-derived optimal correlation threshold that maximized SL recall and enrichment while minimizing the number of screened gene pairs. This cutoff ($r = 0.25$) was translated to the human network using a percentile-matching strategy based on Pearson correlation coefficients. In yeast, $r \geq 0.25$ corresponded to the top 0.075% of all gene–gene correlations (Supplementary Fig. 1C).

The equivalent percentile in the human correlation matrix corresponded to $r \geq 0.13$, preserving comparable network sparsity and resolution (Supplementary Fig. 1D). This translated threshold ($r = 0.13$) was applied in the partial correlation clustering to terminate merging and define the final set of modules.

To further improve SL discovery potential, modules were expanded to include paralogous genes not already present in the modules. Paralogs were identified according to criteria established in the In4mer library design[6]. Specifically, gene pairs were retained if they exhibited high sequence identity and were constitutively expressed across human cell lines, as assessed using Ensembl BioMart[7] and CCLE expression data[8].

<u>Functional enrichment analysis</u>

Functional enrichment of network modules was performed using a one-tailed hypergeometric over-representation test implemented in SciPy[9], with Bonferroni–Holm correction for multiple comparisons. For stability, a small pseudo-count was added before log transformation. Pathway over-representation was quantified both by statistical significance (P-value) and by an effect-size fold change (FC), which compares the proportion of pathway genes present in a module with the proportion expected under a uniform random draw from the background. Pathways were retained if the adjusted $P < 0.01$ and fold-change enrichment > 2 (i.e., $Log_2$ FC > 1).

For the yeast network, enrichment was performed using Gene Ontology (GO) annotations (Ashburner et al., 2000). For the human network, enrichment was evaluated using Reactome and GO (BP, MF, CC) reference libraries obtained from MSigDB[2,10,11]. Pathways containing more than 400 genes were excluded to reduce annotation bias and multiple-testing burden. To integrate both statistical significance and magnitude of effect, a Functional Enrichment Score was computed for each cluster–pathway pair as:

$$Functional\ Enrichment\ Score = -Log_{10}\ Adj\ Pval\ \times\ Log_2 FC$$

This composite metric was used to rank and visualize biologically coherent pathways in both yeast and human networks.

<u>Network modules included in screens</u>

Two major functional modules were selected from the human functional interaction network for experimental screening: the Receptor Tyrosine Kinase (RTK, Fig. 1F) and the DNA Damage Response module (DDR, Fig. 5A). These are large, hierarchically organized clusters each containing multiple smaller functional units linked by shared pathway components and moonlighters. Modules falling outside the size and functional enrichment criteria were generally deprioritized due to lower likelihood of GI enrichment. However, because coessentiality reflects shared dependencies across diverse cancer lineages, biologically related genes do not always co-cluster; highly context-specific, redundant, or lineage-restricted essentiality can partition paralogs or pathway components in separate modules. To capture these relationships and enable cross-module testing, we additionally retained a small set of functionally coherent modules for screening based on their relevance to the primary RTK and DDR clusters (Supplementary Fig. 1F, Fig. 5A).

In addition to the main RTK module, we included a few smaller functionally related modules that connect to RTK activity and ER processing. These cover (i) core glycosylation needed for RTK maturation (FUT8 pathway), (ii) RAS/MAPK regulators and a RAS–RAF initiation set (NF1/SPRED; KRAS/NRAS–RAF1), and (iii) canonical downstream pathways: PI3K–AKT–mTOR and MAPK with their phosphatase controls (BRAF–MAP2K1/2–MAPK1/3, DUSPs) (Supplementary Fig. 1F).

The main DDR module (Fig. 5A, 121 genes) is a large cluster spanning the canonical DNA damage and replication-stress response: ATR–ATRIP checkpoint signaling, fork protection, and γ-H2AX signaling. Repair pathways include homologous recombination (BRCA1/2, RAD51 paralogs) and Fanconi anemia, plus base-excision/SSB repair (XRCC1, APEX2, FEN1), and end-joining (LIG4–XRCC4) with POLQ-mediated alt-EJ. The module also contains replication factors (RFC2–5, POLE3/4) and an iron–sulfur cluster biogenesis arm, supporting DNA metabolism. In addition to the main DDR module, we retained a few small, functionally related sets: a checkpoint/p53 set (ATM–CHEK2–TP53), a p53 ubiquitin-control set (MDM2/MDM4–USP7), a genome-maintenance set (SMC5/6–TOP3A–RMI), and a chromatin module (ATRX–DAXX) (Fig. 5A). These were included to ensure coverage of checkpoint control, p53 regulation, recombination/replication recovery, and genome stability pathways.

### In4mer genetic interaction library design and construction

enAsCas12a guide RNA sequences were selected using the CRISPick algorithm[12,13]. The entire database of target sequences was downloaded and the top four guides ("pick order") for each gene were retained. Nontargeting sequences from Esmaeili Anvar, Lin *et al.*[6] were used. We selected 50 essential and 50 nonessential reference genes from Hart *et al.*[14,15]. A list of 199 target genes from the RTK module and 167 genes from the DDR module were collected, as well as the list of 4,435 paralogs from the Inzolia library[6].

For single genes (n=4,374), two guide arrays targeting each gene were designed. The two clones were designed as in the Inzolia library:

| | | | | |
|---|---|---|---|---|
| Clone_1 | guide 1 | guide 2 | guide 3 | guide 4 |
| Clone_2 | guide 2 | guide 1 | guide 4 | guide 3 |

where guide1 is the guide targeting the gene with pick order 1 from CRISPick.

For RTK, DDR, and paralog genetic interactions, two guide arrays targeting each gene pair were designed. The two clones are:

| | | | | |
|---|---|---|---|---|
| Clone_1 | gene 1_guide 1 | gene 2_guide 1 | gene 1_guide 3 | gene 2_guide 3 |
| Clone_2 | gene 2_guide 2 | gene 1_guide 2 | gene 2_guide 4 | gene 1_guide 4 |

where gene1_guide1 is the guide targeting gene 1 with pick order 1 from CRISPick. The RTK sublibrary has 39,402 guide arrays and the DDR sublibrary has 27,722 guide arrays. All guide arrays targeting paralog pairs from the Inzolia library were also added to the genetic interaction library. Note that, since paralogs are explicitly added to both the RTK and DDR networks, the paralog_pair annotation can duplicate RTK and DDR gene pairs.

### Library production

Library production was performed as previously described[6]. Briefly, oligonucleotide pools were synthesized by Twist Bioscience using the following template:

5′-*[Forward_primer]***cgtctcgAGAT**[sgRNA]TAATTTCTACTATTGTAGAT[sgRNA]AAATTTCTACTCTAGTAGAT[sgRNA]TAATTTCTACTGTCGTAGAT[sgRNA]TTTTTT**GAATggagacg***[Reverse_primer]*-3′

Individual oligonucleotide pools were amplified using the following primer sets and cloned into the pRDA-550 vector (Addgene #203398) using the NEBridge Golden Gate Assembly Kit (BsmBI-v2) (New England Biolabs) according to the manufacturer's protocol. Desalted ligation products were transformed into Endura electrocompetent cells (Lucigen), and plasmid libraries were purified using the PureLink™ Expi Endotoxin-Free Plasmid Purification Kit (Thermo Fisher Scientific). Library quality and representation were assessed by next-generation sequencing to verify uniformity and completeness.

Primer set; Forward primer, 5'-3'; Reverse primer, 5'-3'; Library

1; AGGCACTTGCTCGTACGACG; TGTGGGCCCGGCACCTTAA; Genetic interaction

2; GTGTAACCCGTAGGGCACCT; GTCGAGAGCAGTCCTTCGAC; Negative control

3; CAGCGCCAATGGGCTTTCGA; AGCCGCTTAAGAGCCTGTCG; Validation

### Cell culture and generation of enAsCas12a expression cells

A375, HT29, MDAMB468, HCT116, LS180 and SW48 cell lines were obtained from American Type Culture Collection (CRL-1619, HTB-38, HTB-132, CCL-247, CL-187, and CCL-231). PC9 cell line was obtained from Millipore Sigma (90071810-DNA-5UG). KP4, A549, SNB75 and CORL105 cell lines were gifted from Tim Heffernan. SW837 cells were gifted from Ronald DePinho. HT29, MDAMB468, PC9, SW837, HCT116, SNB75, SW48, and CORL105 cells were routinely cultured in RPMI-1640 supplemented with 10% fetal bovine serum, 1 mM sodium pyruvate and 2 mM L-glutamine. A375 and A549 cells were routinely cultured in DMEM supplemented with 10% fetal bovine serum, 1 mM sodium pyruvate and 2 mM L-glutamine. KP4 and LS180 cells were routinely cultured in RPMI-1640 supplemented with 20% fetal bovine serum, 1 mM sodium pyruvate and 2 mM L-glutamine. All cell lines were authenticated by short tandem repeat (STR) profiling performed by the Cytogenetic and Cell Authentication Core Facility at The University of Texas MD Anderson Cancer Center. Cells were routinely tested for mycoplasma contamination using the MycoAlert Mycoplasma Detection Assay (Lonza).

Cells were maintained at 37°C in a humidified incubator with 5% $CO_2$ and passaged regularly to sustain exponential growth. An additional copy of enCas12a was introduced into all cell lines prior to screening via the lentiviral vector pRDA-174 (Addgene #136476), as previously described[16]. Cells expressing enAsCas12a were selected using 10 μg/ml blasticidin.

**Lentivirus production and enCas12a screens**

Lentivirus was generated by the University of Michigan Vector Core. Although virus stocks were not pre-titered, multiplicity-of-infection (MOI) tests were performed for each cell line to determine the optimal infection conditions yielding a 30-50% infection rate[17]. Based on library size, transductions were performed as previously described[6], ensuring that at least 1,000 cells per sgRNA were maintained after puromycin selection. Polybrene and puromycin were applied at the following concentrations: polybrene was used at 1 μg/ml for A375 and 8 μg/ml for all other cell lines; puromycin was used at 2 μg/ml for most cell lines, with the exceptions of A375 (1 μg/ml) and MDAMB468 (0.75 μg/ml).

Following selection (confirmed by 0% viability in non-transduced control cells after puromycin treatment), surviving cells were pooled and divided into two replicates. For each replicate, at least 500x library coverage was maintained throughout the screen. At the screen endpoint (7-10 doublings post-transduction), cells were harvested at least 500x coverage per replicate for genomic DNA isolation.

**Genomic DNA extraction and sequencing**

Genomic DNA (gDNA) was extracted using the Mag-Bind Blood & Tissue DNA HDQ Kit (Omega Bio-tek) and quantified with a Qubit fluorometer (Thermo Fisher Scientific). Illumina-compatible guide array amplicons were generated directly from purified gDNA by a one-step PCR. In detail, indexed PCR primers were synthesized by Integrated DNA Technologies (IDT) and incorporated standard 8-nt Illumina index sequences (D501-D508 and D701-D712)[6]. To maintain adequate library representation, at least 200X gDNA coverage was amplified per replicate. Each gDNA sample was divided into multiple 50 μL reactions, each containing up to 2.5 μg gDNA. Reaction composition was as follows: 1 μL each primer (10 μM), 1 μL 50X dNTP mix, 5% DMSO, 5 μL 10X Titanium Taq Buffer, and 1 μL 10X Titanium Taq DNA Polymerase (Takara). PCR conditions were: initial denaturation at 95 °C for 60 s; 25 cycles of 95 °C for 30 s and 68 °C for 1 min; final extension at 68 °C for 3 min.

Following PCR amplification, reactions from each sample were pooled and purified by E-Gel™ SizeSelect™ II Agarose Gels, 2% (Thermo Fisher Scientific). Purified amplicons were quantified by Qubit and assessed by D1000 ScreenTape Assay (Agilent), yielding expected fragment sizes of ~360 bp for the Genetic interaction library and ~372 bp for the Negative control and the Validation libraries.

Amplicons were pooled and supplemented with 30% of a custom random library amplicon to increase sequence diversity. Sequencing was performed on an Illumina NextSeq 2000 platform using forward custom primers (synthesized by IDT)[6]. The Genetic interaction library was sequenced using a 151-8-8 single-end read format, while the Negative control and Validation libraries were sequenced using a 158-8-8 single-end read format.

### Screen analysis

Data processing and normalization

The sequencing reads were mapped to the library using only perfect matches. Raw read count data from each sample were first processed and normalized. Arrays with fewer than 10% of the median read count in the plasmid library (90 reads) were excluded. After this filtering, genes and gene pairs represented by fewer than 2 remaining arrays were also removed. A pseudocount of one was added to all remaining counts, and all read counts were converted to read frequency by dividing by the total mapped counts in the sample. Guide-level $\log_2$ fold change (FC) was calculated as the log ratio of read frequency at the endpoint relative to $T_0$. Gene-level LFCs were then derived by averaging the LFCs of all guides targeting the same gene, and replicate means were computed.

LFCs were then Z-transformed into Zfc. For each cell line, the distribution of LFCs was fitted to a two-component normal distribution using the Gaussian Mixture function in *scikit-learn* (sklearn.mixture). The component with the higher weight represented the null bulk of genes and gene pairs with no effect on fitness, while the component with the lower weight, smaller mean, and larger variance captured genes and gene pairs whose knockout impaired cell growth. The mean and standard deviation of the higher-weight distribution were used to calculate the Zfc as follows:

$$Zfc_{i,j} = \frac{FC_{i,j} - \mu_{high\,weight,j}}{\sigma_{high\,weight,j}}$$

where *i* represents all target single genes and gene pairs, and *j* represents the 12 cell lines.

Quality control and cell line selection

The library included 50 essential and 50 nonessential genes as positive and negative controls, respectively, selected from the Hart reference sets[14,15]. Fold change distributions for essential and nonessential controls are shown in Supplementary Fig. 3. Screen quality was assessed using Cohen's *D*, defined as the difference in mean LFC between nonessential and essential controls divided by the pooled standard deviation:

$$Quality\ Score = Cohen's\ D = \frac{mean\ LFC_{nonessential} - mean\ LFC_{essential}}{pooled\ standard\ deviation}$$

We observed a clear bimodal distribution of Cohen's D values, with eight high and four others substantially lower. The eight cell lines with the highest *Cohen's D* scores—MDAMB468, PC9, A375, HT29, HCT116, KP4, LS180, and SW837—were retained for subsequent analyses.

GRAPE: Genetic interaction Regression Analysis of Pairwise Effects

Genetic interaction (GI) scores were calculated using a regression-based model to estimate single-gene fitness effects. The predictor was a binary matrix *A(i,j)*, where rows (*i*) represented individual arrays and columns (*j*) represented target genes across all pairwise arrays. Each matrix element was assigned *A(i,j) = 1* if array *i* included a perturbation of gene *j*, and 0 otherwise. Because single-gene arrays contained four guides per gene, while dual-gene arrays contained two guides per gene, the corresponding entries for single-gene arrays were manually set to 2 to account for an empirically observed increase in fitness defect when targeting a gene with four guides vs. two. The response vector was the observed, normalized gene-level Zfc. Regression coefficients ($\beta$) obtained from the model represented the inferred single-gene knockout fitness effects from the model (Fig. 2A).

Using these coefficients, we predicted the expected additive LFC for each gene pair (Fig. 2B and Supplementary Fig. 4). Gene pairs with predicted LFC below the minimum observed LFC were excluded to maintain predictions within the assay's dynamic range. The raw GI score was then defined as the difference between observed and predicted LFC, capturing deviations from additivity.

To correct for heteroscedasticity – increased variance at lower predicted LFC – a sliding window normalization was applied. For each gene pair, the local standard deviation was estimated from the 500 nearest neighbors ranked by predicted FC (Fig. 2C). Zgi were calculated as the raw GI divided by the local standard deviation (Fig. 2D; Supplementary Fig. 6). Statistical significance of synergistic (negative GI) and suppressive (positive GI) interactions was then determined using p-values from the N(0,1) distribution, followed by multiple testing corrections using the Benjamini–Hochberg method to obtain adjusted p-values /false discovery rates for final interaction calls.

Importantly, the GRAPE regression coefficient estimates of single gene knockout fitness are driven by the observed phenotype from the pairwise knockout arrays for each sublibrary (~400 arrays for each gene in RTK and ~350 in DDR genes), outweighing the contribution from the dedicated single gene knockout arrays (two for each gene). We measured the single gene beta coefficients vs. the observed single gene knockout (Supplementary Fig. 5) and note that there is high concordance between beta coefficients and observed SKO fold changes. One consistent outlier, *POLE*, was removed from further analysis.

<u>Calculating effect size (ES)</u>

The effect size of a genetic interaction is the fraction of the maximum possible effect of the pairwise knockout, where an effect size of 1 implies that the maximum observable (negative) fold change was measured, and an effect size of zero implies that the observed fold change is exactly the expected fold change as calculated by GRAPE coefficients. To measure maximum possible effect size, the read count of guide arrays targeting each gene pair was set to zero across all replicates in all screens, and raw fold change was calculated while holding every other read count unchanged. Maximum fold changes per gene pair target were averaged across replicates and guide arrays and transformed to Zfc using the same null bulk mean and standard deviation learned above (see *Data processing and normalization*). In principle, the max observable Zfc calculated here should approach the observed Zfc for guides with near zero reads, but experimental artifacts (e.g. when sequencing depth greatly exceeds library coverage) can put an artificial floor under observed Zfc. We normalized max observable FC to the $10^{th}$-ranked most severe FC observed in each screen, adding a constant to the max Zfc; the results of the scaling are shown in Supplementary Figure 7. Effect size was then calculated as:

$$ES = \frac{GI\_raw}{Max\ observable\ effect} = \frac{(observed\ Zfc - expected\ Zfc)}{(max\ Zfc - expected\ Zfc)}$$

where expected Zfc is a GRAPE output. Effect sizes > 1.0, for the top 10 ranked observed Zfc, are clipped at 1.0. When observed Zfc approaches maximum Zfc, effect size approaches 1.0, and where observed Zfc approaches expected Zfc, effect size approaches zero. Note that all these calculations are based on negative genetic interactions/synthetic lethality. When observed Zfc is greater than expected Zfc, genetic interactions are positive/suppressive and effect size becomes negative. When expected Zfc is very negative, the denominator can become quite small, and large (<< -1) negative effect sizes can result.

Classifying synthetic interactions

For each cell line, high-effect-size (HE) synthetic interactions were defined by the criteria Zgi Padj < 0.25, Zfc < -2.0, and ES > 0.15, where Zgi, Zfc, and ES are described above. HE positive interactions were, similarly, Padj (suppressor) < 0.25, Zfc > -1, and ES < -0.15.

Integrated Z score, Zint

To more sensitively recover interaction signals across all screens, we sought to combine individual screen-level Z scores for each gene pair into an integrated Z-score. The naïve approach, Stouffer's method:

$$Z_{Stouffer} = \frac{\sum Zgi}{\sqrt{n}}$$

is N(0,1) distribution under the null hypothesis only if the per-screen Z-scores are mutually independent. This is not the case; the Z-scores from the twelve screens are significantly correlated, with a mean off-diagonal of the correlation matrix from the full dataset of 0.425. However, the full-data correlation matrix is biased toward more positive values by signal-driven covariance from the minority of pairs with real, shared genetic interactions across cells. To remove this bias, we estimated the covariance $\Sigma\_null$ across the null bulk of the pooled RTK + DDR matrix by considering only rows where max(|Zgi|) < 3, censoring ~9% of all gene pairs. This still resulted in a mean off-diagonal correlation of 0.358. Given this degree of correlation, uncorrected Stouffer was inflated by:

| **Panel** | **Subset** | **Stouffer SD** | **corrected SD** | **inflation** |
|---|---|---|---|---|
| DDR | HiQ | 2.515 | 1.349 | **1.86**× |
| DDR | all_12 | 2.921 | 1.344 | **2.17**× |
| RTK | HiQ | 2.036 | 1.089 | **1.87**× |
| RTK | all_12 | 2.408 | 1.100 | **2.19**× |

Without correction, every Z would be inflated roughly two-fold and the false-discovery rate at any nominal cutoff would be very anti-conservative.

To correct for correlated samples, we used Strube's method:

$$Z_{int} = Z_{Strube} = \frac{w' \, Zgi}{\sqrt{w' \, \Sigma\_null \ w}}$$

wherein $w$ is a vector of sample weights, and $\Sigma\_null$ is the null bulk covariance matrix described above. We used equal weights, since using Cohen's D as a weight vector yielded nearly identical results and the range of weights is modest. Applying the method on the negative control library using $\Sigma\_null$ estimated from its own null bulk yielded a distribution of Zint scores with standard deviation = 1.00, as expected, and with the three positive control paralog synthetic lethals showing extreme scores of Zint < -8.6.

Within- and between-module GI enrichment

We quantified enrichment of genetic interactions both within and between modules in the human network using only assayed gene–gene pairs and treating pairs as unordered. For each interaction class, synthetic lethal and suppressor, we first computed the global hit rate across the screened universe to serve as the background. Within-module enrichment was calculated as the $\log_2$ ratio of the observed hit frequency among screened pairs whose genes are both in the module to the global background rate. Between-module enrichment was calculated similarly, using only screened cross-pairs whose genes lie in the non-overlapping portions of two modules (to respect hierarchy and avoid double counting shared genes). Enrichment values were reported as $\log_2$(Observed/Expected) frequency. Analyses were performed separately for SL and suppressor GI, as shown in Supplementary Figures 12-13.

**Validation studies**

Competitive growth assay

To validate the relative fitness effects of candidate hits identified from RTK screens, sgRNAs targeting either non-targeting controls or the corresponding genes (Supplementary table S16) from hits were cloned into the pMV-AA-131 vector (gift from John Doench), which co-expresses mCherry and a puromycin resistance marker. For each construct, lentivirus was individually produced and used to transduce HT29 and PC9 cells. Following transduction, cells were selected with puromycin for 3 days to enrich for mCherry positive populations. After completion of selection, the mCherry-positive cells were mixed with pre-sorted GFP-expressing corresponding control cells at a 1:4 ratio. The mixed populations were plated and imaged using the Incucyte S3 live-cell analysis system. Images were acquired every 12 hours for seven days using the green and red fluorescence channels, with exposure times of 300 ms and 400 ms, respectively. To assess relative fitness over time, fluorescence intensity ratios (green/red) from later time points were normalized to the baseline ratio measured 24 hours post-seeding.

Validation library

We designed a small, all-by-all library highly enriched for interactions discovered in our screens but small enough to be applied to *in vivo* models. We selected 25 genes with strong interactions, primarily from the transport and glycosylation network in Figure 3F, and added STT3A/B and

SOS1/2 as positive controls for a total of 29 genes. A library targeting all pairs of the 29 genes was designed using the In4mer constructs as previously described, with three constructs per target:

| | | | | |
|---|---|---|---|---|
| Clone_1 | gene 1_guide 1 | gene 2_guide 1 | gene 1_guide 2 | gene 2_guide 2 |
| Clone_2 | gene 2_guide 3 | gene 1_guide 3 | gene 2_guide 1 | gene 1_guide 1 |
| Clone_3 | gene 1_guide 2 | gene 2_guide 2 | gene 1_guide 3 | gene 2_guide 3 |

for gene pairs, and:

| | | | | |
|---|---|---|---|---|
| Clone_1 | gene 1_guide 1 | non-targeting | gene 1_guide 2 | non-targeting |
| Clone_2 | non-targeting | gene 1_guide 3 | non-targeting | gene 1_guide 1 |
| Clone_3 | gene 1_guide 2 | non-targeting | gene 1_guide 3 | non-targeting |

for single knockouts. Including 50 reference essentials, the total library size was 1,455 guide arrays.

<u>3D validation using pancreatic cancer organoid models</u>

Pancreatic patient-derived organoid PDO065[18] (*KRAS*$^{G12V}$) was cultured in DMEM/F12 (WNT-conditioned media-1:1) growth media supplemented with 10 mM Nicotinamide (HY-B015, MCE), 1 mM N-Acetylcysteine (MCE), 10 nM Gastrin (MCE), B27 (Gibco), 1 mM HEPES, Glutamax (1X), 0.5 µM A83-01 (MCE), 10 µM SB-202190 (MCE), 50 ng/mL EGF (R&D), Pen/Strep, and Glutamine (1X). During routine passage, culture media were supplemented with 10 µM Y-27632 (Tocris) and 2.5 µM CHIR 99021 (Tocris). Lentivirus generated from the validation library were transduced to PDOs by spinoculation at 600xg for 1 hr, in the presence of 10 µM Y-27632 and 8 µg/mL polybrene. Organoids were then embedded into Matrigel (Corning) domes 24 hours post-transduction. Y-27632 was removed with the beginning of puromycin selection three days after lentivirus transduction. After untransduced cells were eliminated, transduced organoids were split into replicates, maintained at 500x, and passaged weekly for 8 doublings post-transduction. Prior to harvesting, organoids were digested with TrypLE, neutralized with FBS, counted, and washed with PBS. Genomic DNA was extracted using the DNeasy Blood & Tissue kit (Qiagen) and processed following the same sample preparation protocol used for 2D screening.

<u>In vivo validation using colorectal cancer xenograft models</u>

All xenograft studies were conducted in accordance with institutional ethical guidelines and were approved by the University of Texas MD Anderson Cancer Center Institutional Animal Care and Use Committee (IACUC). B1003 is a *BRAF*$^{V600E}$ patient-derived colorectal cancer xenograft model as previously described[19,20]. B1003 PDX cells were routinely cultured in RPMI-1640 supplemented with 10% fetal bovine serum, 1 mM sodium pyruvate and 2 mM L-glutamine. HT29 cells and B1003 PDX were transduced at a low multiplicity of infection (MOI $\approx$ 0.3) using lentivirus generated from the validation library. Transduced cells were selected with puromycin (2 µg/mL) for three days. On day 4 post-transduction, $2 \times 10^6$ cells were injected subcutaneously into the flanks of 8-week-old female athymic nude mice (Envigo) (n = 3 mice per line). Tumor growth was monitored using digital calipers, and tumor volume was calculated as (length $\times$ width$^2$)/2. Tumors were harvested once the average tumor volume reached approximately 500 mm$^3$. gDNA was extracted from whole tumor tissues using the Mag-Bind Blood & Tissue DNA HDQ Kit according to the manufacturer's instructions. Library preparation and sequencing were performed following the same protocol used for the in vitro (2D) screens, with a minimum of 50 PCR reactions (~125 µg gDNA) per sample.

<u>Analysis of validation screens</u>

Recall and precision were calculated using positive and negative reference sets derived from the data. For positives (true synthetic lethal interactions), we identified 22 interactions with Zint FDR < 1%, where each interaction scores as a high-effect-size hit in at least 1 screen. These interactions included the *STT3A/B* and *SOS1/2* positive controls. For reference noninteracting pairs, we selected 65 gene pairs from the validation library with Zint score > 1 in the 8 cell line screens.

**DDR screen comparison**

To benchmark our DDR library, we compared it against three published DNA damage response (DDR) genetic interaction datasets: Fielden *et al.*[21], Hayward *et al.*[22], and Herken *et al.*[23], using data obtained from the Supplementary Information or source materials of each study. We first assessed overlap at both the single-gene and gene-pair levels across the four libraries, identifying 21 shared single genes and 210 shared gene pairs (Fig. 6B).

We next compared synthetic interaction hits among these 210 common gene pairs. Hit thresholds were defined according to each study's reported criteria: adjusted p-value for the Zint score < 0.1 for our screen and for Hayward *et al.*; Gemini score ≤ –1 for Fielden et al.; and GI score < –3.46 for Herken *et al.* Applying these thresholds identified 3 hits in our screen, 13 in Fielden *et al.*, 10 in Hayward *et al.*, and 23 in Herken et al., with no gene pairs scoring as hits across all four studies. To visualize the extent of overlap among hits, we generated an UpSet plot summarizing hit concordance across the four studies (Fig. 6C).

**Estimating background GI frequency with a negative control library**

<u>Negative control library design</u>

To establish a baseline for genetic interaction frequency and evaluate enrichment of predicted SL modules, we designed a combinatorial CRISPR library composed of randomly paired genes with minimal prior evidence of interaction. The library was comprised of 64 genes spanning a range of essentiality and expression levels, but very low coessentiality scores. Three synthetic lethal paralog pairs (CNOT7–CNOT8, HDAC1–HDAC2, PITPNA–PITPNB) were added as positive controls, with each paralog being paired with all other genes in the library, for a total of 70 genes. To minimize the inclusion of biologically connected pairs, any gene pairs with known interactions in BioGRID[24] were avoided. Constructs were designed used a three array per target format, as shown here:

for single-gene:

| | | | | |
|---|---|---|---|---|
| Clone_1 | guide 1 | non-targeting | guide 2 | non-targeting |
| Clone_2 | non-targeting | guide 1 | non-targeting | guide 3 |

| | | | | |
|---|---|---|---|---|
| Clone_3 | guide 2 | non-targeting | guide 3 | non-targeting |

for dual-gene:

| | | | | |
|---|---|---|---|---|
| Clone_1 | gene 1_guide 1 | gene 2_guide 1 | gene 1_guide 2 | gene 2_guide 2 |
| Clone_2 | gene 2_guide 1 | gene 1_guide 1 | gene 2_guide 3 | gene 1_guide 3 |
| Clone_3 | gene 1_guide 2 | gene 2_guide 2 | gene 1_guide 3 | gene 2_guide 3 |

The library was cloned and constructed as described for other libraries, and screened in three cell lines using standard protocols, with samples collected on day 14. After sequencing, guide arrays with read counts < 3 in the plasmid pool were censored (7% of the pool, which otherwise had median read count 1,178, and 92% of guide arrays with >50 reads), and then both single genes and gene pairs were censored if they did not have three guide arrays. After filtering, all 70 single genes and 1,915 gene pairs remained, including all six paralogs and all three paralog pairs (Supplementary Fig. 9).

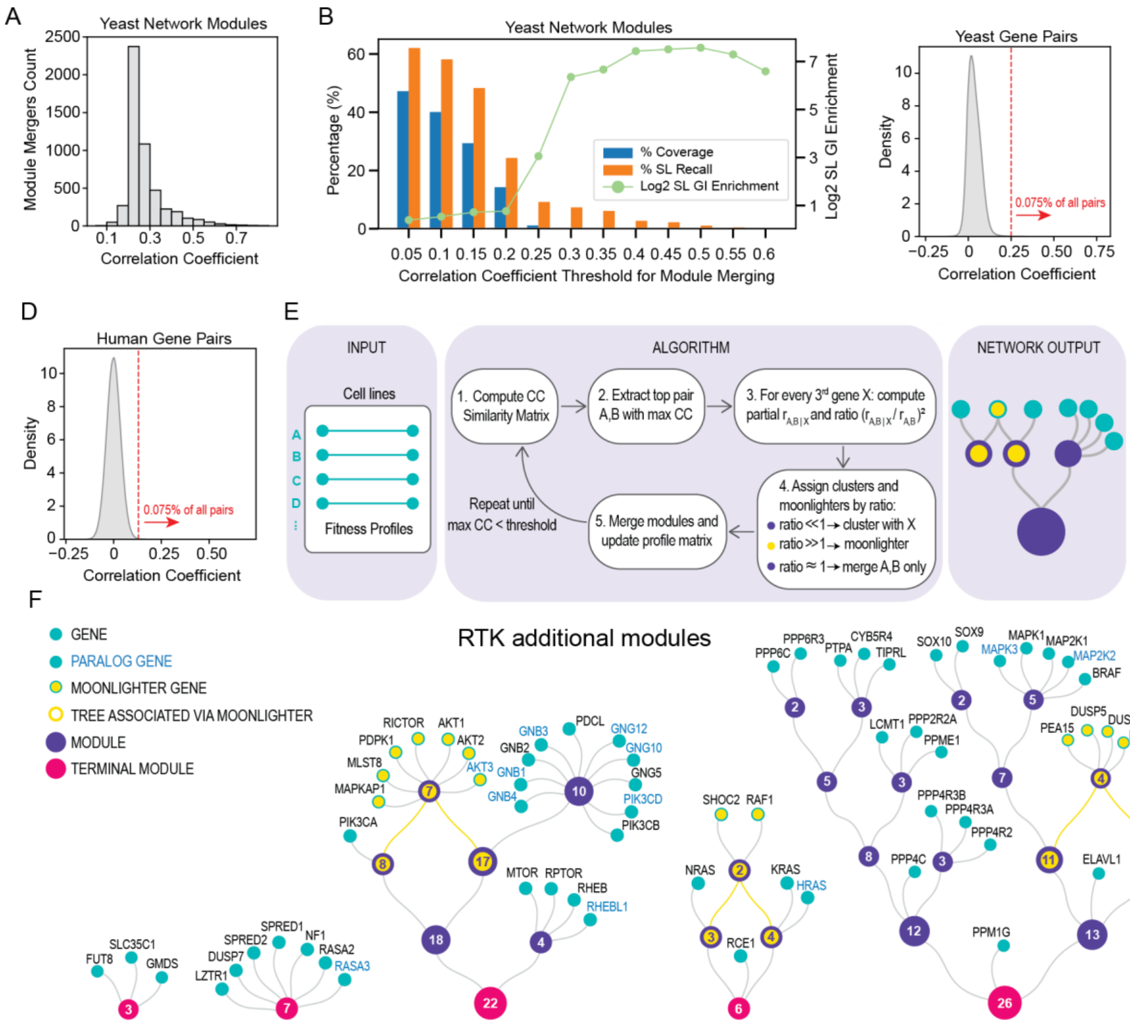


**Fig. S1. Yeast-to-human genetic interaction prediction model using functional networks.**

(A) Distribution of module merger events vs. correlation coefficient thresholds, as measured by partial correlation–based clustering of yeast data. (B) Trade-off between SL recall, coverage, and Log fold SL enrichment across correlation thresholds. Bar plots show the percentage of screened gene pairs (blue) and percentage of SL recall (orange) at each CC cutoff. The green line indicates $\log_2$ fold SL enrichment within modules at each cutoff. (C-D) Translation of the yeast correlation threshold to the human network via percentile matching. Density plots of covariance-whitened Pearson correlation coefficients for all gene pairs in the yeast (C) and human (D) networks, with the matched percentile thresholds indicated by vertical red dashed lines. (E) Schematic of the partial correlation method for clustering of functional networks. (F) Additional RTK modules that were manually included in screening.

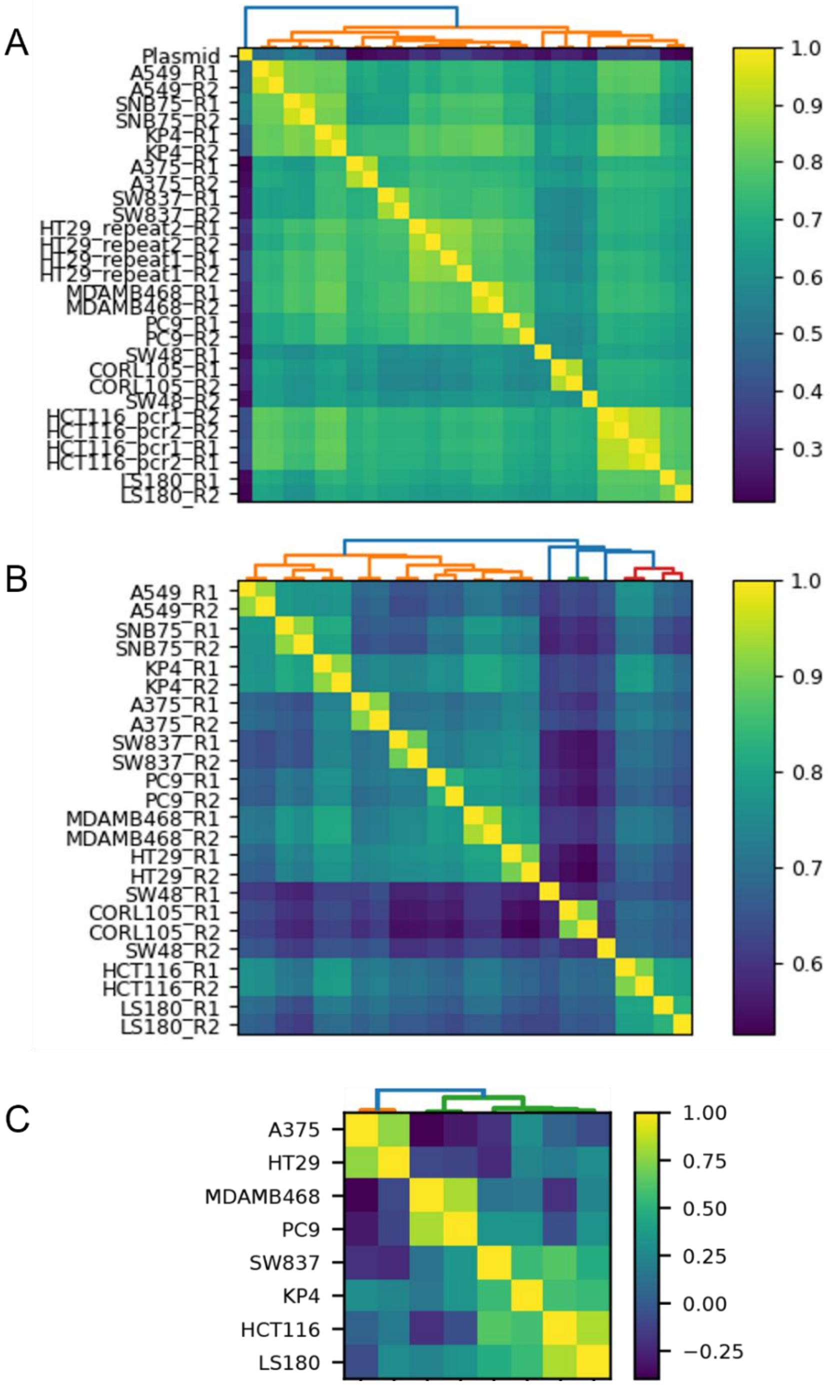


**Fig. S2. Quality control of genetic interaction library.** (A) Clustermap of raw read counts across all samples from all 12 cell lines. HT29 repeat 1 and repeat 2 are full biological replicate screens. HCT116 pcr 1 and pcr 2 are replicate amplicon PCR from the same genomic DNA extraction. (B) Clustermap of $\log_2$ fold-change across all 24 replicates before averaging. (C) Targeted clustermap of 13 RTK genes in 8 HiQ cell lines. Colorbars: Pearson correlation coefficient.

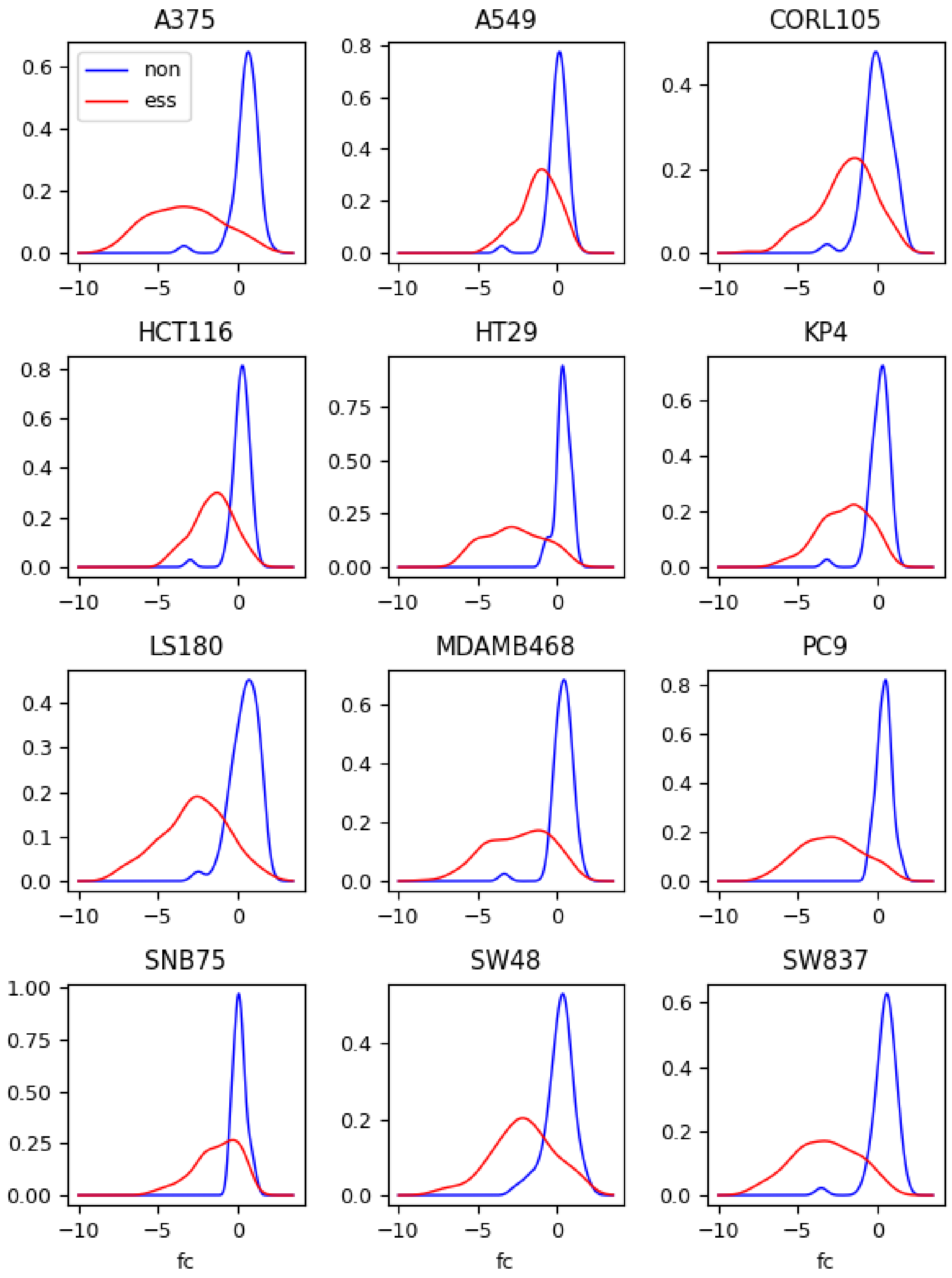


**Fig. S3. Calculation of Cohen's D for quality control.** Kernel density estimate of fold changes for guide arrays targeting reference essential (red) and non-essential (blue) controls across 12 genetic interaction screens. Mean of replicates and mean of guide arrays targeting each gene/gene pair. Cohen's D is measured from these data and is used as a metric of screen quality.

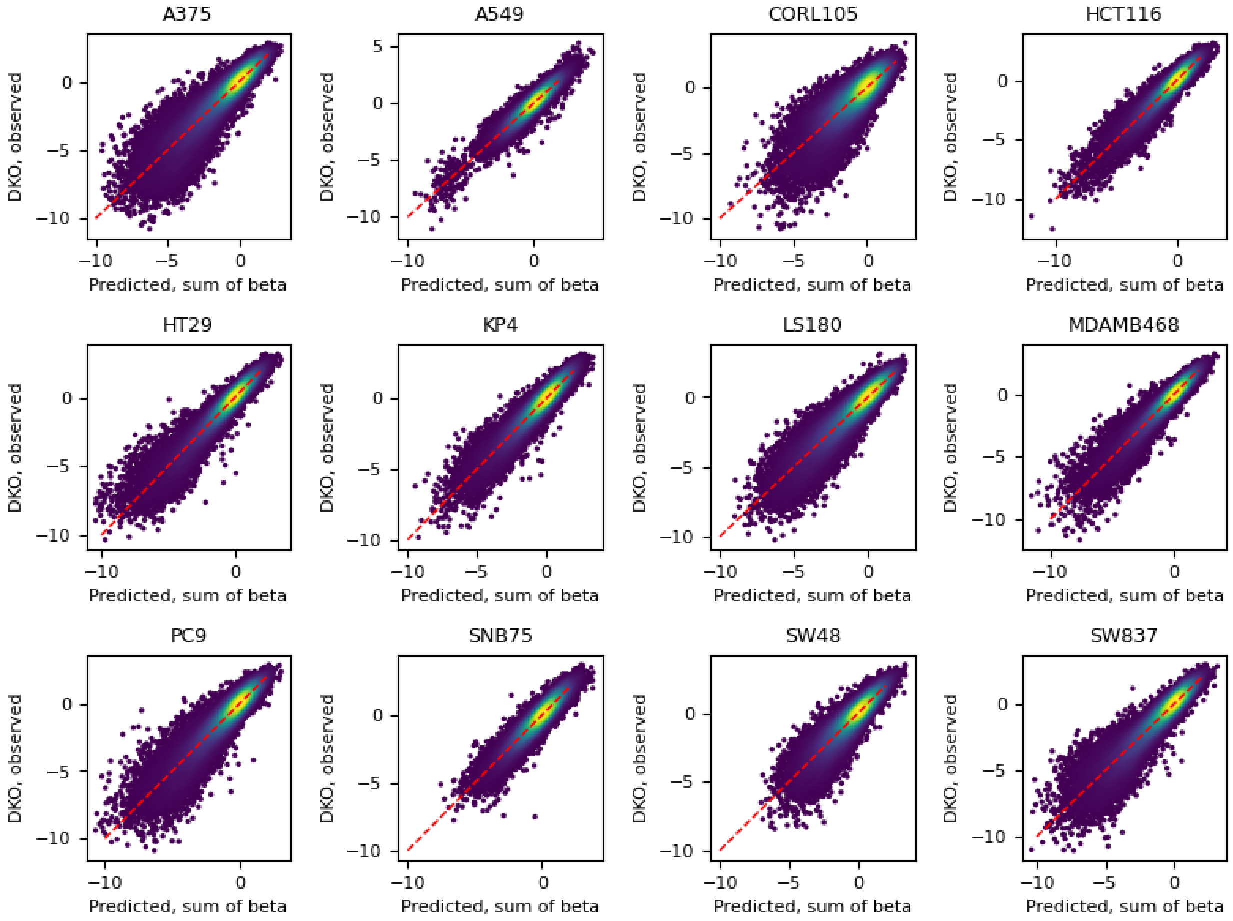


**Fig. S4. GRAPE quality control.** Scatter plot of observed (y-axis) versus predicted (x-axis) Z-normalized fold-change for all gene pairs in 12 genetic interaction screens. Predicted fold change for each gene pair is the sum of the beta coefficients for the two genes as learned by GRAPE.

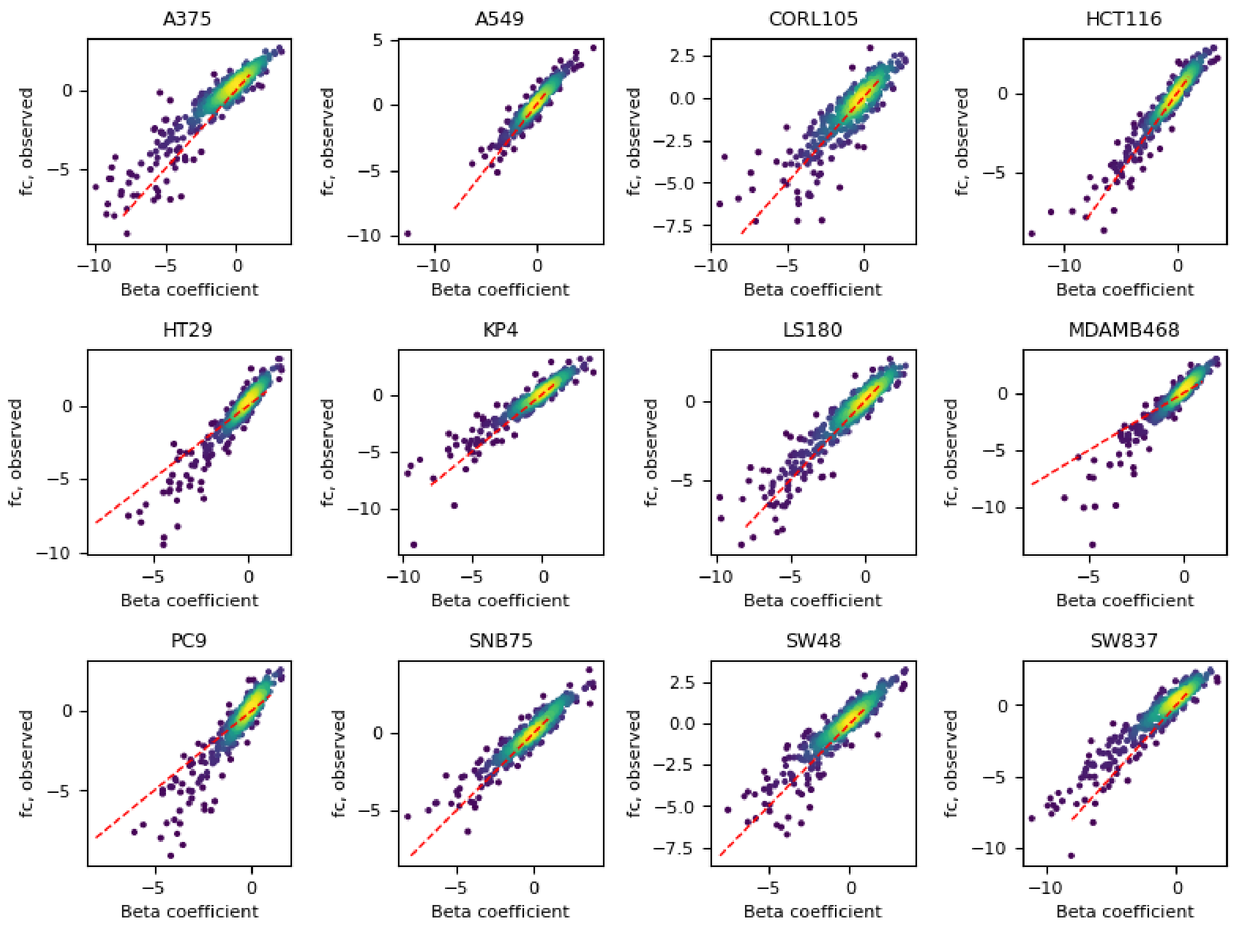


**Fig. S5. Scatter plots of observed versus predicted fold-change for all single-gene knockouts in 12 genetic interaction screens.** Predicted fold change is the single gene beta coefficient from GRAPE. Observed is the fold change from guide arrays targeting single genes. The strong outlier *POLE* displayed extreme deviation from the regression model and was removed for following analysis.

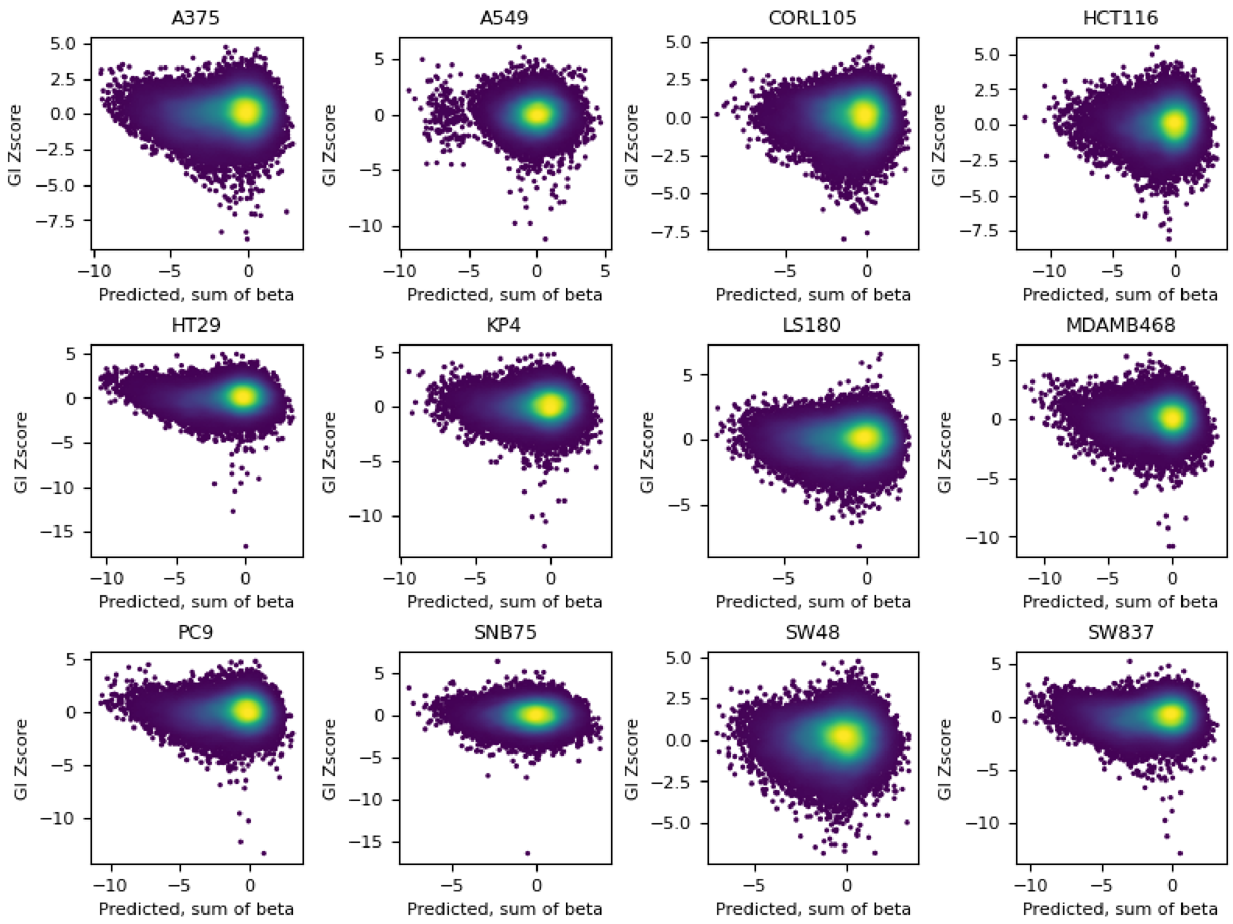


**Fig. S6. Scatter plots of GI Z score from GRAPE versus predicted fold-change for all pair knockouts.** Negative GI z scores reflect synthetic-lethal effects.

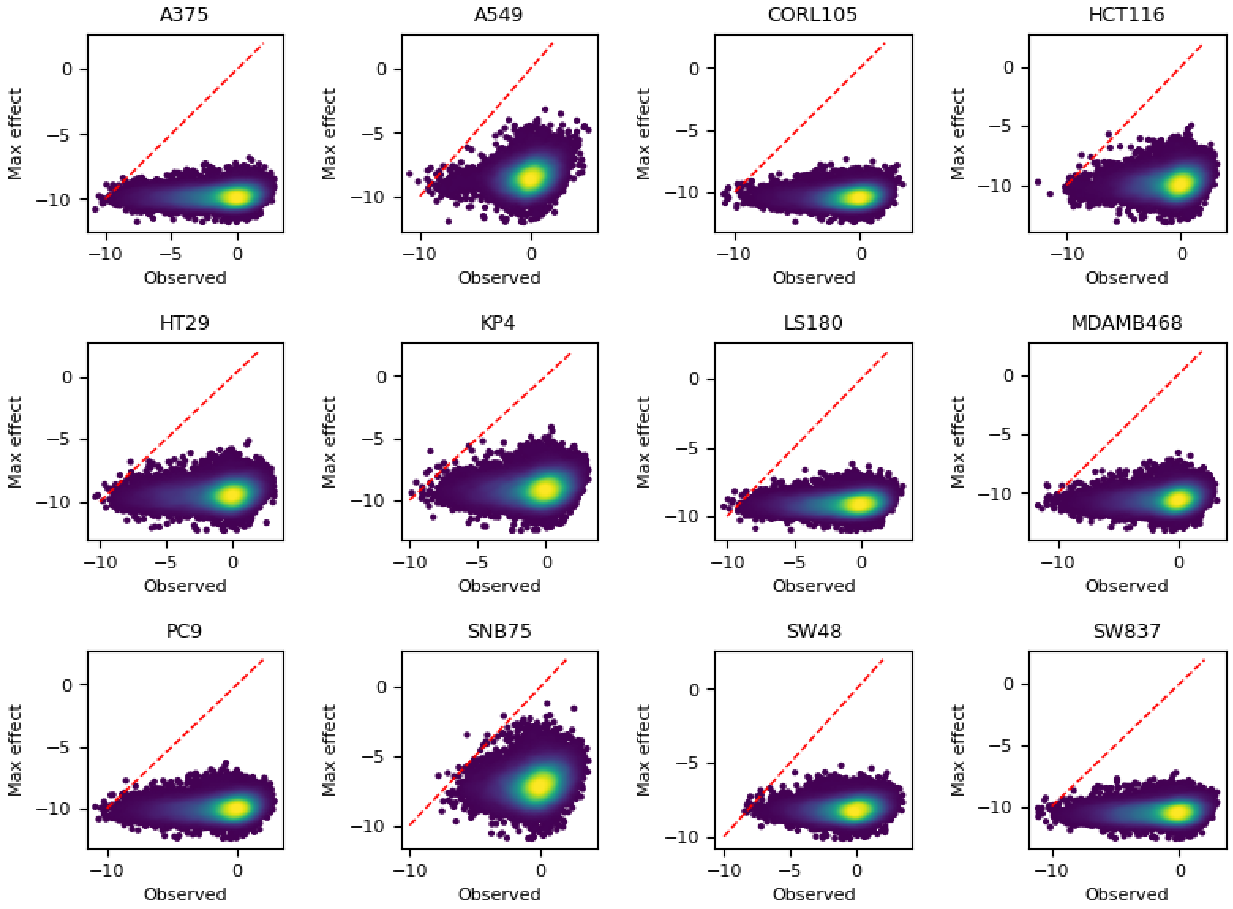


**Fig. S7. Calculation of effect size.** For each target, the observed fold change (x-axis) is plotted against the maximum observable fold change (y-axis), if all guide arrays targeting the gene/genepair yield zero reads in all replicates. The ratio of observed to max fold change is the effect size.

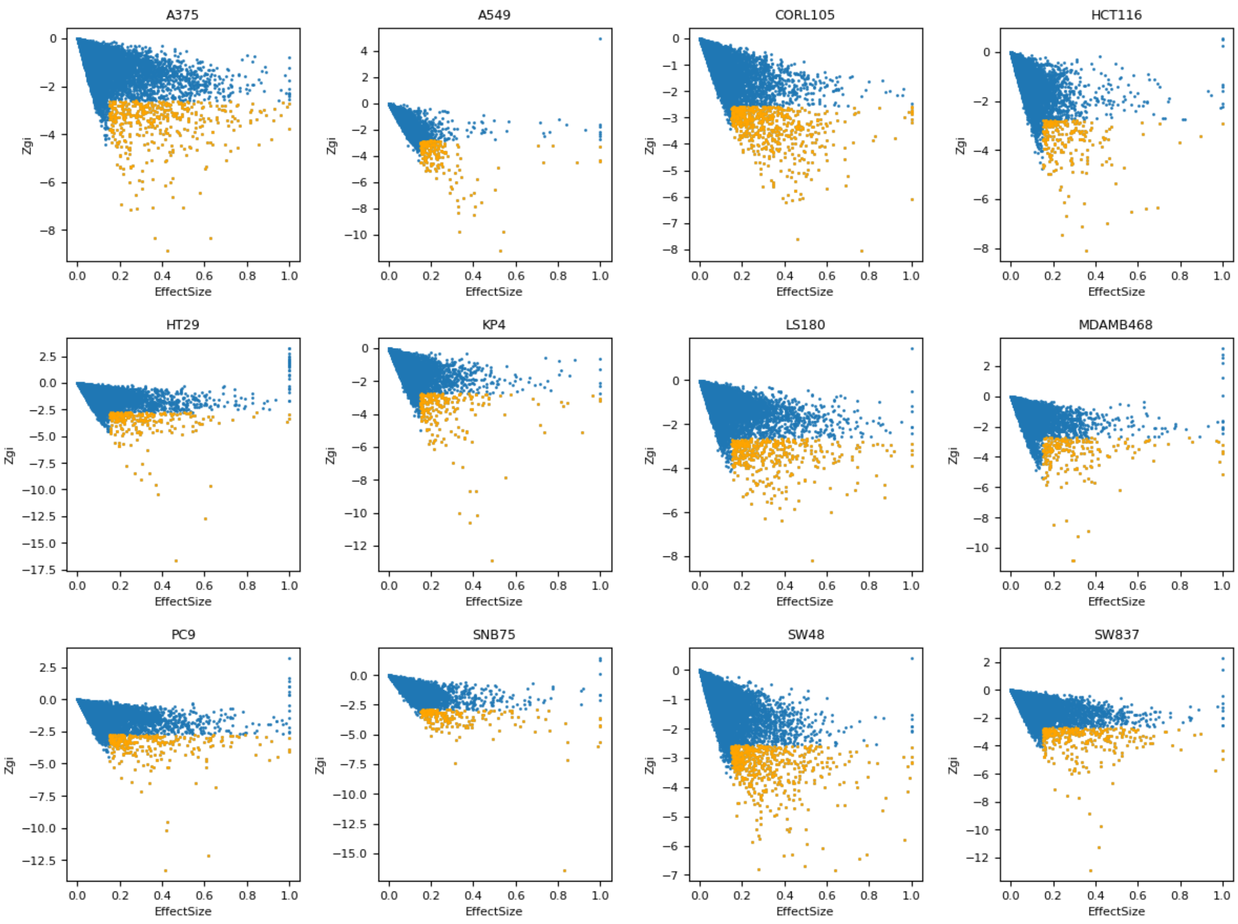


**Fig. S8. Effect size vs. Zgi.** For each screen, positive effect size (synthetic lethals) is plotted against Zgi score. Yellow, targets with Effect Size > 0.15 and Zgi FDR < 0.25.

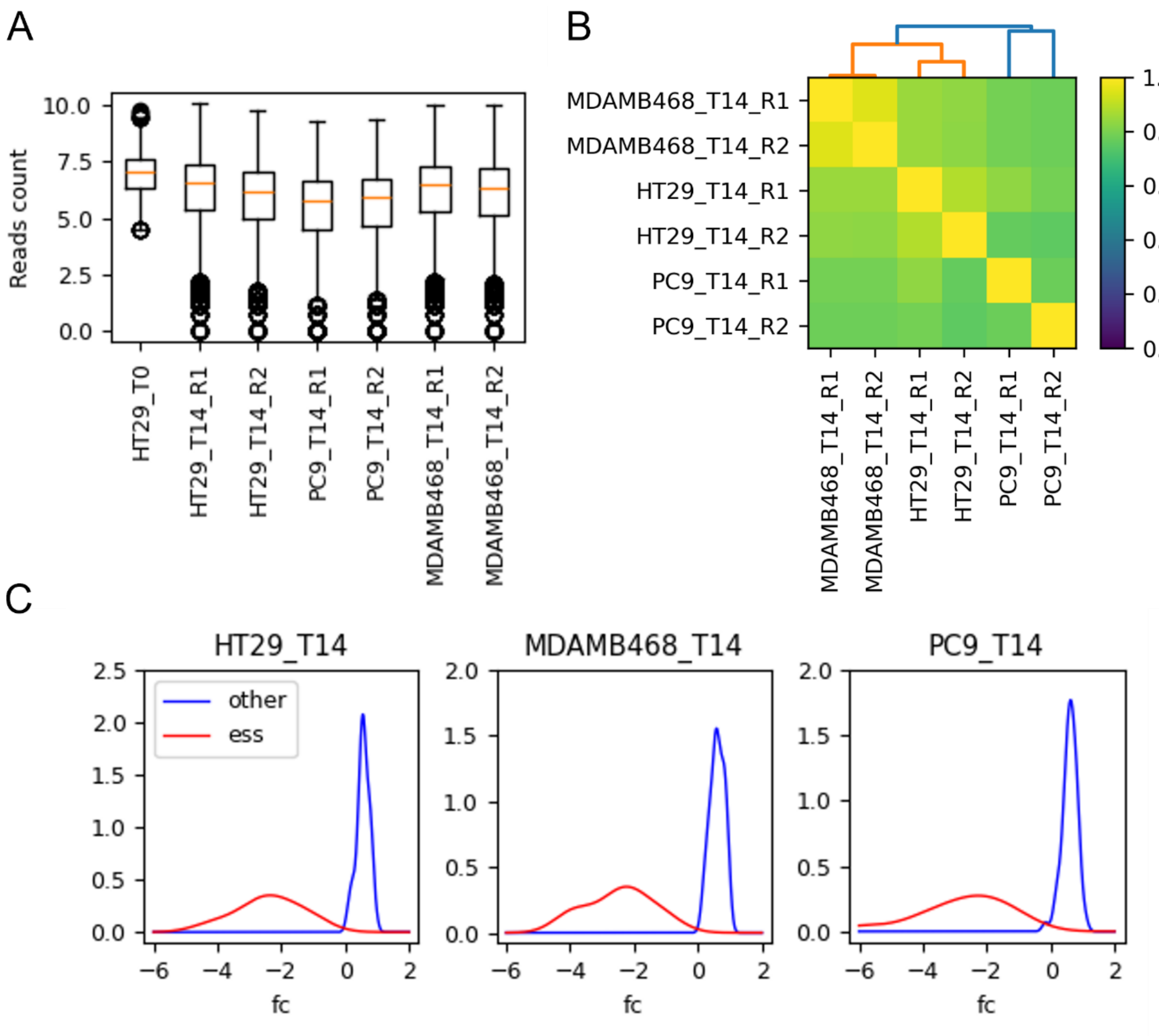


**Fig. S9. Quality control of Negative control library screens.** (A) Raw read-count distributions for all samples. (B) Clustermap of $\log_2$ fold-change patterns across three cell lines. Colorbar indicates Pearson correlation coefficient. (C) Kernel density estimates of $\log_2$ fold-change distributions of guide arrays targeting reference essential controls (red) and other targets (blue) controls in NC screens.

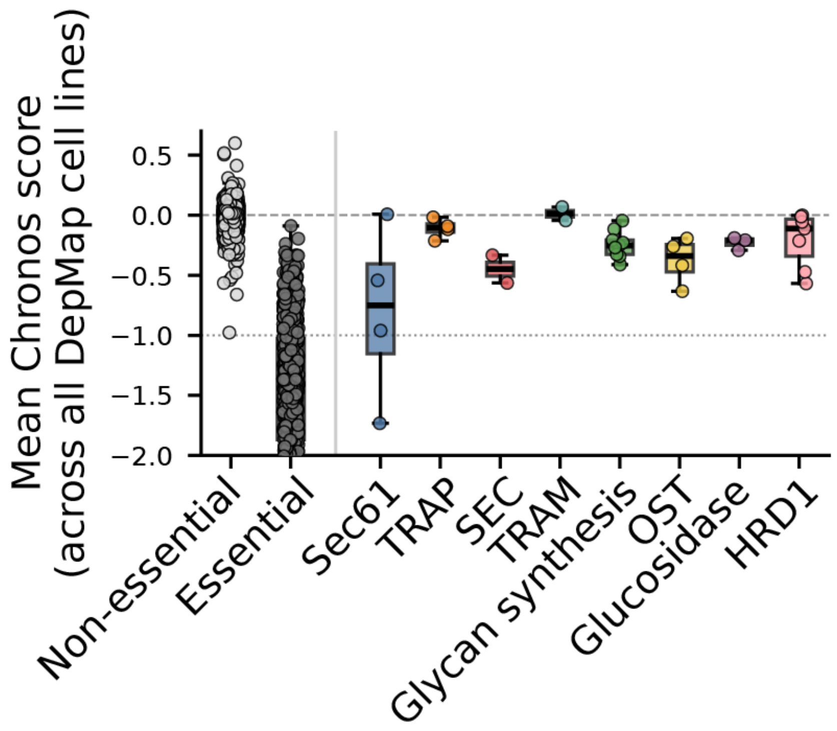


**Fig. S10. Gene essentiality of glycosylation module.** Mean Chronos score across DepMap of genes in each protein complex, compared to reference essentials and nonessentials.

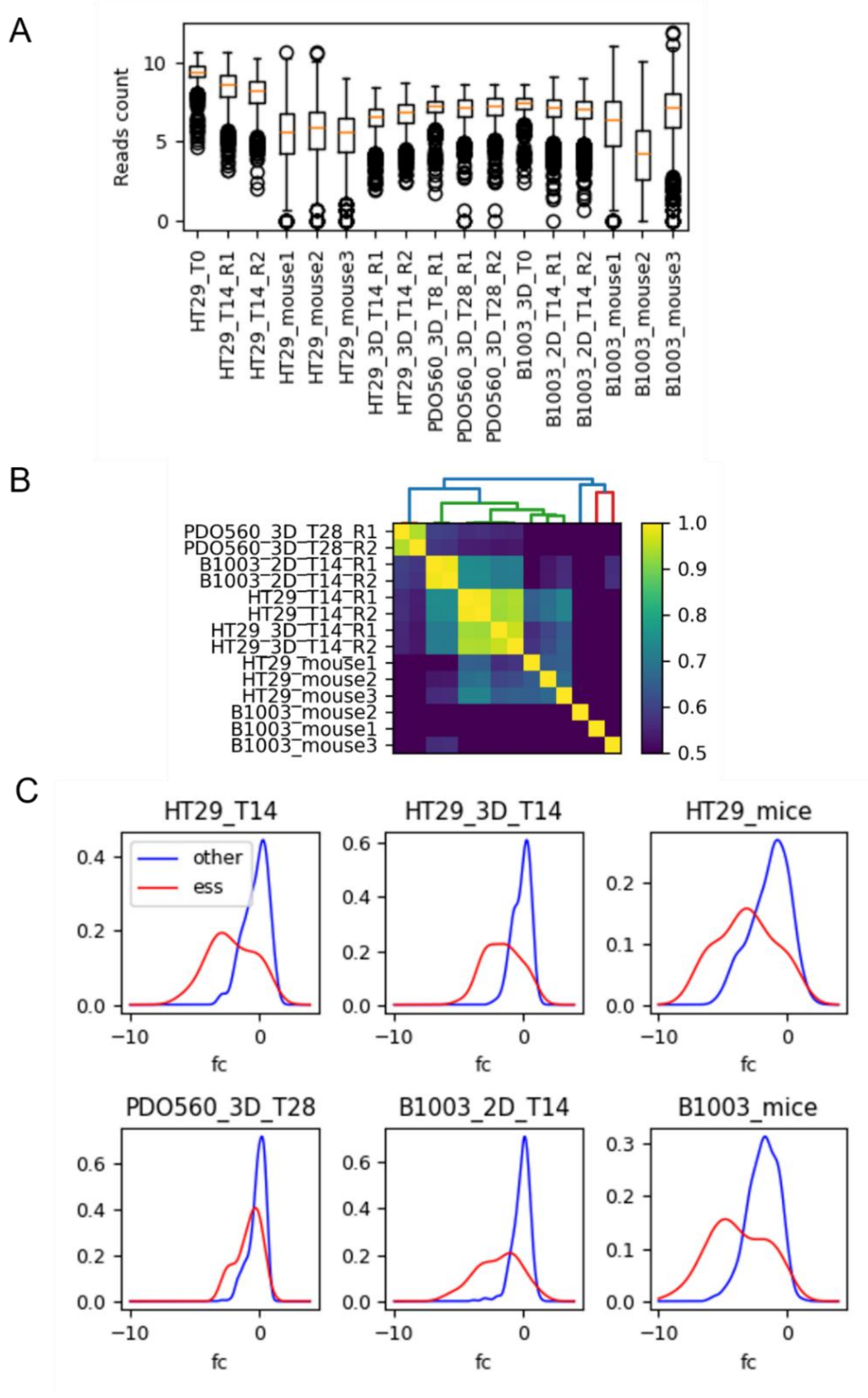


**Fig. S11. Quality control of validation library screens performed in 2D, 3D, and PDX models.** (A) Boxplots of log2 read-counts for all samples. Guides with fewer than 90 reads in the HT29 T0 reference sample (log2(90) = 6.5) were excluded from downstream analysis. (B) Clustermap of $\log_2$ fold-change across all models. Colorbar indicates Pearson correlation coefficient. (C) Kernel density estimates of fold-change distributions of guide arrays targeting reference essential controls(red) and other targets(blue) in validation screens.

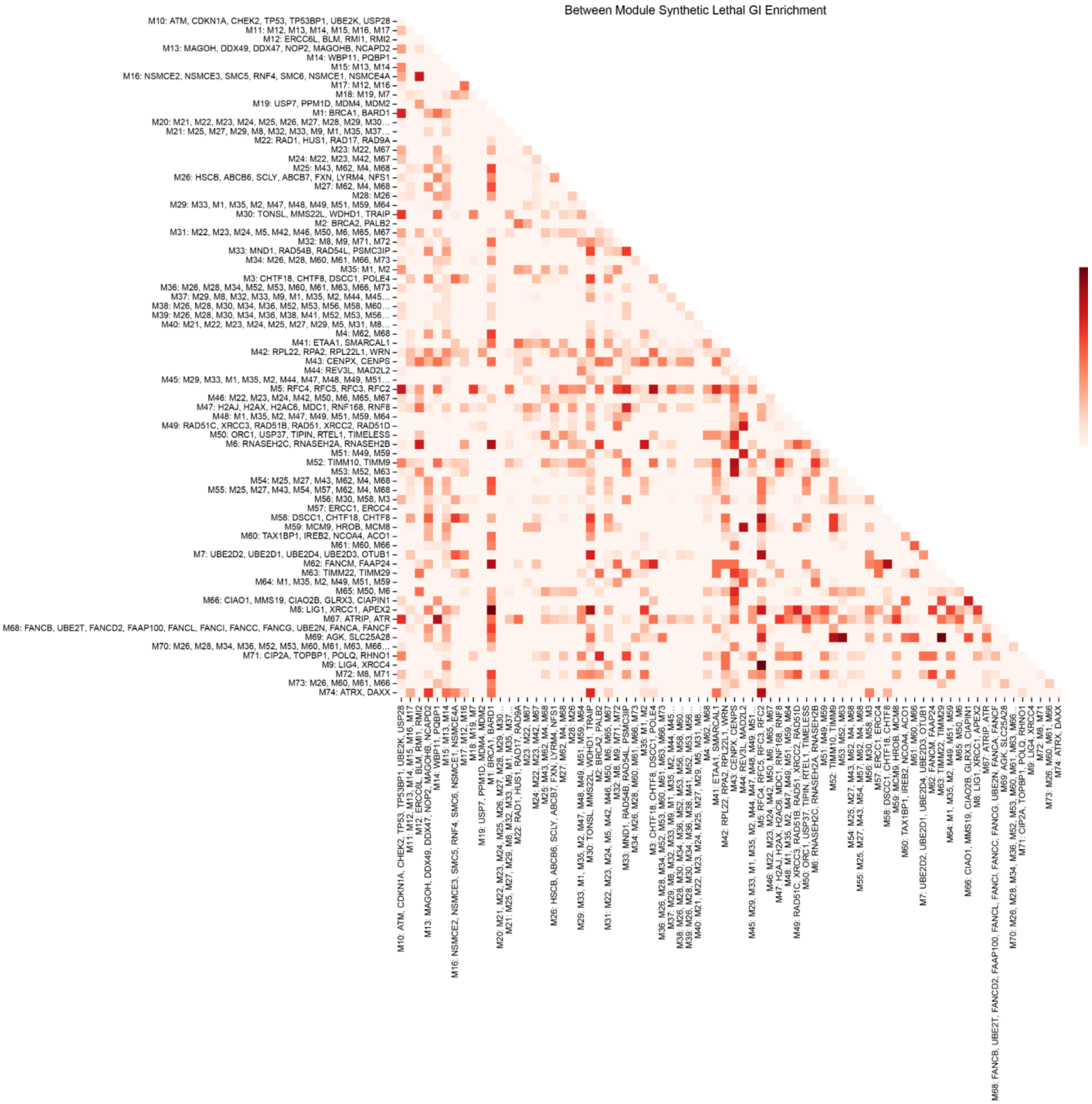


**Fig. S12. Enrichment of SL within and between DDR network modules.** Labels show gene names for the first-level nodes in the hierarchy; higher-level merge nodes display the merged module IDs (capped at 10). Color intensity is based on log2(observed/expected) frequency for synthetic lethal enrichments between modules. Diagonal indicates within-module values.

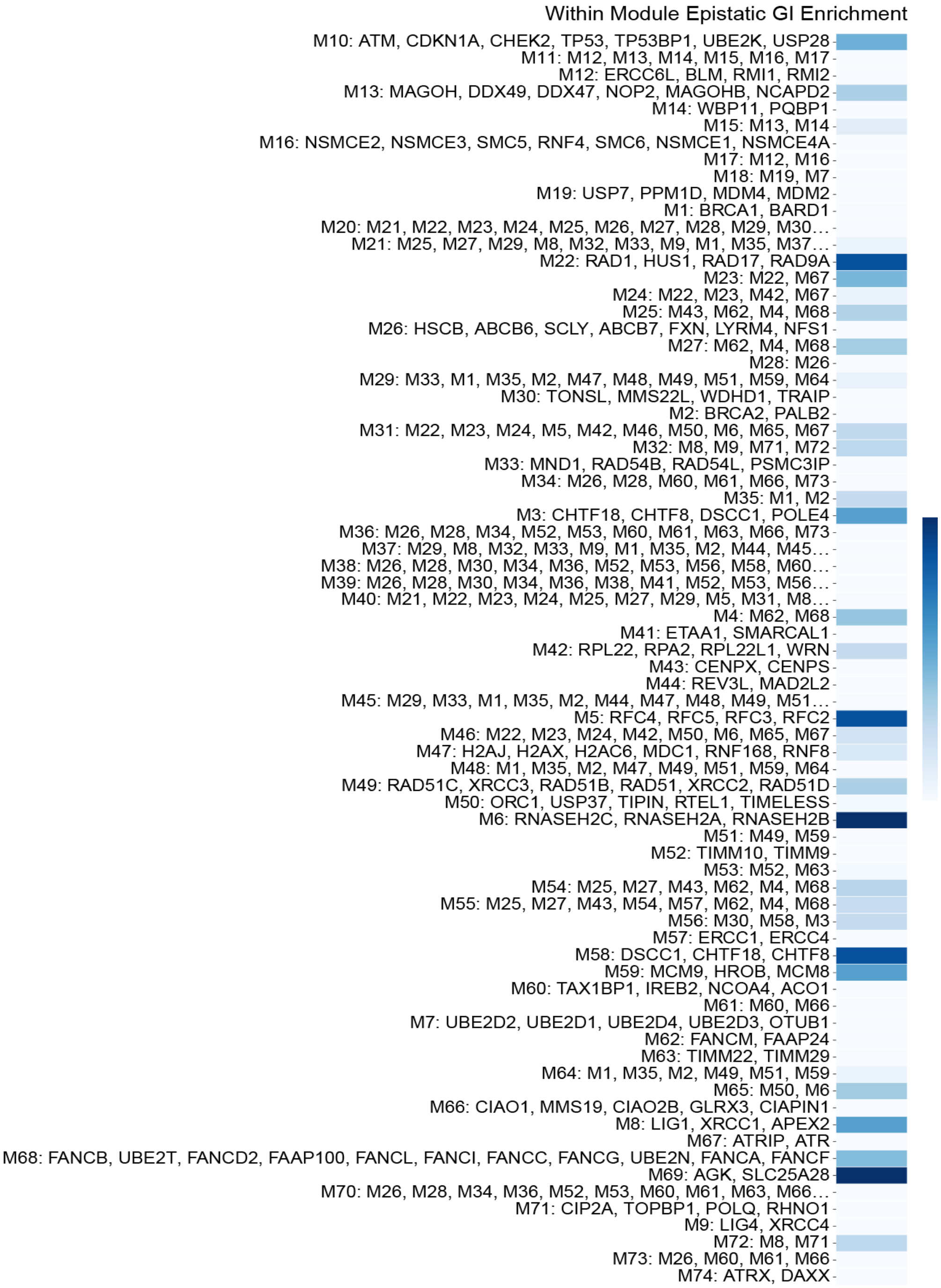


**Fig. S13. Within-module epistatic GI enrichment for DDR network modules.** Labels show gene names for first-level nodes; higher-level merge nodes list merged module IDs (capped at 10). Color encodes $\log_2$(observed/expected) frequency for within-module suppressor interactions.
